# A novel implementation of spinal fMRI demonstrates segmental organisation of functionally connected networks in the cervical spinal cord: A test-retest reliability study

**DOI:** 10.1101/2023.02.27.530185

**Authors:** Olivia S. Kowalczyk, Sonia Medina, Dimitra Tsivaka, Stephen B. McMahon, Steven C. R. Williams, Jonathan C. W. Brooks, David J. Lythgoe, Matthew A. Howard

**Affiliations:** Department of Neuroimaging, Institute of Psychology, Psychiatry & Neuroscience, King’s College London, London, UK; The Wellcome Centre for Human Neuroimaging, Queen Square Institute of Neurology, University College London, London, UK; Medical Physics Department, Medical School, University of Thessaly, Larisa, Greece; Wolfson Centre for Age Related Diseases, King’s College London, London, UK; School of Psychology, University of East Anglia, Norwich, UK

**Keywords:** Spinal fMRI, resting-state fMRI, reliability, test-retest

## Abstract

Resting fMRI studies have identified intrinsic spinal cord activity, which forms organised motor (ventral) and sensory (dorsal) resting-state networks. However, to facilitate the use of spinal fMRI in, for example, clinical studies, it is crucial to first assess the reliability of the method, particularly given the unique anatomical, physiological, and methodological challenges associated with acquiring the data. Here we demonstrate a novel implementation for acquiring BOLD-sensitive resting-state spinal fMRI, which was used to characterise functional connectivity relationships in the cervical cord and assess their test-retest reliability in 23 young healthy volunteers. Resting-state networks were estimated in two ways: (1) by extracting the mean timeseries from anatomically constrained seed masks and estimating voxelwise connectivity maps and (2) by calculating seed-to-seed correlations between extracted mean timeseries. Seed regions corresponded to the four grey matter horns (ventral/dorsal and left/right) of C5-C8 segmental levels. Test-retest reliability was assessed using the intraclass correlation coefficient (ICC) in the following ways: for each voxel in the cervical spine; each voxel within an activated cluster; the mean signal as a summary estimate within an activated cluster; and correlation strength in the seed-to-seed analysis. Spatial overlap of clusters derived from voxelwise analysis between sessions was examined using Dice coefficients. Following voxelwise analysis, we observed distinct unilateral dorsal and ventral organisation of cervical spinal resting-state networks that was largely confined in the rostro-caudal extent to each spinal segmental level, with more sparse connections observed between segments (Bonferroni corrected *p* < 0.003, threshold-free cluster enhancement with 5000 permutations). Additionally, strongest correlations were observed between within-segment ipsilateral dorso-ventral connections, followed by within-segment dorso-dorsal and ventro-ventral connections. Test-retest reliability of these networks was mixed. Reliability was poor when assessed on a voxelwise level, with more promising indications of reliability when examining the average signal within clusters. Reliability of correlation strength between seeds was highly variable, with highest reliability achieved in ipsilateral dorso-ventral and dorso-dorsal/ventro-ventral connectivity. However, the spatial overlap of networks between sessions was excellent. We demonstrate that while test-retest reliability of cervical spinal resting-state networks is mixed, their spatial extent is similar across sessions, suggesting that these networks are characterised by a consistent spatial representation over time.

## 1 Introduction

Spinal cord functional magnetic resonance imaging (fMRI) is a novel but rapidly developing field (Kinany, Pirondini, Micera, et al., 2022; Powers et al., 2018). Combined with brain fMRI, it holds promise for investigation of information processing across all levels of the central nervous system in both health and disease.

Like the brain, the spinal cord is characterised by spontaneous fluctuations in the blood-oxygen-level-dependent (BOLD) signal in the absence of overt stimulation. This intrinsic activity of the spinal cord has been shown to form organised resting-state networks, which can be broadly divided into motor and sensory (Harrison et al., 2021). Reports of strong temporal correlations between the sensory (dorsal) horns and motor (ventral) horns within the cervical spinal cord have dominated the spinal fMRI resting-state literature (Barry et al., 2014, 2016; Eippert et al., 2017; San Emeterio Nateras et al., 2016; Weber et al., 2018). Furthermore, unilateral sensory networks have also been observed in resting spinal data, which were imited in rostro-caudal extent, corresponding to the underlying segmental anatomy of the cord (Kong et al., 2014). Early evidence from simultaneous brain-spine fMRI has also shown that spinal and cerebral resting-state networks are correlated, suggesting a unified functional architecture of intrinsic networks in the central nervous system (Vahdat et al., 2020).

Brain resting-state fMRI is frequently used as a biomarker for identification of neurodivergent states/conditions or treatment effects (Drysdale et al., 2017; Pfannmöller & Lotze, 2019; Taylor et al., 2021). Reliable detection of resting-state networks in the spine would extend this approach to information processing occurring at the level of the cord, such as early modulation of noxious signals or motor functioning (Kinany, Pirondini, Micera, et al., 2022; Tinnermann et al., 2021). Acquiring fMRI recordings from the spinal cord, however, faces unique anatomical, physiological, and methodological challenges, including, among others, the small size of the cord, influence of physiological noise, and reliable static magnetic field shimming (Kinany, Pirondini, Micera, et al., 2022; Tinnermann et al., 2021). These challenges can limit the quality of obtained data and thus pose a threat to the reliability of spinal fMRI. To date, the few studies that investigated the reliability of resting-state spinal cord fMRI showed *good* test-retest reliability (intraclass correlation coefficient (ICC) = 0.64-0.7) in network properties using graph theory measures at 3 T (Liu et al., 2016) and *fair* reliability (ICC = 0.54-0.56) in region-to-region connections at 7 T (Barry et al., 2016). A recent assessment of reliability of region-to-region connections at 3 T has further shown that reliability was *fair* to *good* for dorso-dorsal and ventro-ventral connections but *poor* for within and between-hemicord connections across the cervical cord and generally *poor* for all connections within individual segmental levels (Kaptan et al., 2022). These studies, however, assessed test-retest reliability within the same scanning session. Given that longer lag between scans is associated with poorer reliability in cerebral fMRI (Bennett & Miller, 2010, 2013) and that the scanning set up for spinal cord fMRI is considerably more complicated than for cerebral fMRI (Kinany, Pirondini, Micera, et al., 2022; Powers et al., 2018; Tinnermann et al., 2021), investigations of test-retest reliability of spinal cord fMRI that span separate scanning sessions are warranted. Such investigations will indicate the feasibility of using spinal cord fMRI to reliably detect the effects of experimental manipulation or clinical interventions across different visits, such as perturbations related to experimental pain, persistent pain (e.g., postsurgical), or treatment effects.

Test-retest reliability is inherently tied to data quality. Acquiring good quality spinal cord fMRI recordings is complicated by the influences of baseline physiology and susceptibility artefacts related to differing magnetic susceptibility profiles of surrounding tissues (Kinany, Pirondini, Micera, et al., 2022; Saritas et al., 2014; Tinnermann et al., 2021). Shimming procedures can minimise the effects of these factors by reducing magnetic field inhomogeneities. A combination of high order and *z*-shimming is frequently used in spinal cord fMRI to improve signal quality (Eippert et al., 2017; Finsterbusch et al., 2012; Kinany, Pirondini, Mattera, et al., 2022; Vahdat et al., 2020). Nonetheless, while *z*-shimming offers large signal gains by accounting for the off-resonance variation along the cord, implementing simultaneous *x, y*, and *z*-shimming can achieve additional benefits by preventing signal loss caused by magnetic field gradients in left/right and anterior/posterior directions (Islam et al., 2019). Furthermore, given that magnetic field inhomogeneities can induce artefacts in traditional echo planar imaging (EPI) sequences incorporating fat saturation pulses, using a spectral-spatial pulse exciting only tissue water could further improve signal quality (Bernstein et al., 2004).

This study assesses the test-retest reliability of cervical spinal cord resting-state fMRI over two separate scanning sessions. Additionally, we demonstrate a novel implementation for acquiring BOLD-sensitive resting-state spinal fMRI and characterise functional connectivity relationships in the cervical cord in healthy adult volunteers. The acquisition sequence used here operates on a General Electric (GE) scanner platform, using high order shimming and *x, y*, and *z* slice-specific linear shimming, together with spectral-spatial excitation pulses designed to excite tissue water only. This approach reduces signal dropout and increases temporal signal-to-noise ratio (tSNR) within the cervical spinal cord (see Tsivaka et al., In prep for full details of the acquisition method).

### Our pre-registered hypotheses (Kowalczyk et al., 2021) are

1. Discrete resting-state sensory and motor networks should be observable in regions of the dorsal and ventral cervical spinal cord, respectively, using T2*-weighted BOLD EPI.
2. Spinal responses observed during the assessments of hypothesis 1 will be reliable, with ICC inter-session test-retest reliability statistics greater than 0.4.

## 2 Material and methods

### 2.1 Participants

Data from twenty-three healthy right-handed (as assessed by the Edinburgh Handedness Inventory (Oldfield, 1971)) adult volunteers (13 females, mean + SD age: 23.91 ± 3.84 years) were collected for all study visits and survived all quality assessments. Full details of participant/data exclusion are shown in Figure 1.

**Figure 1.**
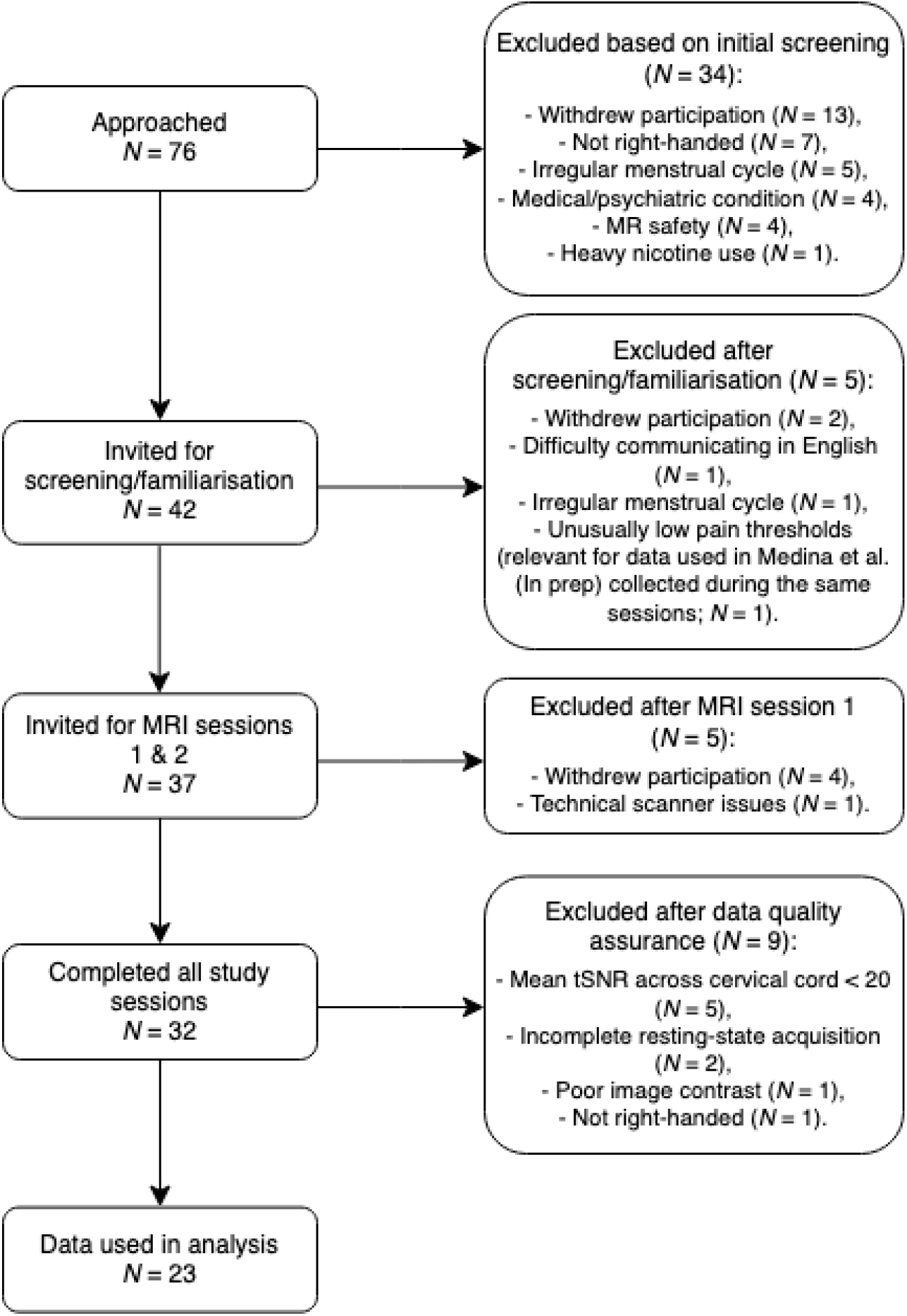
A diagram showing the selection of participants fulfilling the study eligibility criteria and data quality assurance.MR = Magnetic Resonance, tSNR = Temporal Signal to Noise Ratio.

Full inclusion and exclusion criteria for this study are outlined in the study preregistration (Kowalczyk et al., 2021). Briefly, participants were excluded due to: (1) history of psychiatric, medical, or psychological conditions, (2) history of substance or alcohol abuse, (3) regular use of medications affecting the central nervous system, (4) irregular menstrual cycle for female participants, (5) MRI-related contraindications. Additionally, participants were excluded if they were unwilling to adhere to the following lifestyle guidelines before each visit: (1) abstain from alcohol for 24 hours, (2) limit caffeine consumption to one caffeinated drink on each study day, (3) abstain from non-steroidal anti-inflammatory drugs (NSAIDs) or paracetamol for 12 hours, (4) abstain from nicotine-containing products for 4 hours.

Written informed consent was obtained. This study was approved by the Psychiatry, Nursing, and Midwifery Research Ethics subcommittee at King’s College London, UK (HR-16/17-4769).

### 2.2 Procedure

This study comprised three visits – a screening/familiarisation visit and two identical MRI visits for test-retest purposes. The mean (± SD, range) interval between each study visit was 21 (± 22, 1-84) days. Additional measures not described here pertaining to pain modulation and guided motor action were collected during the study visits, see the preregistration (Kowalczyk et al., 2021) and Medina et al. (In prep) for details.

#### 2.2.1 Session 0 – screening and familiarisation

Compliance with study lifestyle guidelines (see Section 2.1) was assessed at the beginning of the session. Participants underwent breath alcohol and urine drugs of abuse tests to check alcohol/substance use. Caffeine, nicotine, and NSAIDs/paracetamol intake were assessed by self-report. Participants were familiarised with the scanner environment by visiting a mock scanner.

#### 2.2.2 Session 1 and 2 – MRI scanning

Sessions 1 and 2 were identical. The sessions began with an assessment of compliance with the study lifestyle guidelines as described above. Additionally, participants completed the state version of the State Trait Anxiety Inventory (STAI) (Spielberger et al., 1971) to assess differences in anxiety levels between sessions. No differences were observed (*t*(22) = 1.23, *p* = 0.223, *d* = 6.12, 95% CI [−0.67; 1.6]; session 1 mean ± SD = 27.61 ± 1.32; session 2 mean ± SD = 29.17 ± 1.49). Subsequently, following optimisation of static 0^th^, 1^st^, and 2^nd^ order shims and linear slice-specific shims, and structural data acquisition (see Section 2.3), a 10 min 50 s resting-state scan was acquired. Participants were instructed to keep their eyes open and look at the fixation cross displayed in the centre of the screen (white cross on a black background). Respiratory and cardiac traces were recorded with respiratory bellows and a pulse oximeter respectively, along with scanner triggers (at the start of each TR), throughout the scan.

### 2.3 MRI acquisition

Data were acquired using a 3T GE MR750 System (General Electric, Chicago, Illinois) equipped with both a 12-channel head, neck, and spine coil and a 4-channel neurovascular array at the NIHR Wellcome King’s Clinical Research Facility, King’s College London. A sagittal 3D CUBE T2-weighted structural image was acquired at the beginning of the scanning session over 64 slices with a coverage of the whole brain and cervical spine to vertebral level T1 (repetition time (TR) = 2.5 s, echo time (TE) = 120 ms, echo train length = 78, flip angle = 90°, field of view (FOV) = 300 mm, acquisition matrix = 320×320, slice thickness = 0.8 mm. This acquisition was based on Cohen-Adad et al. (2021) with the FOV increased to 300mm.

Functional data were acquired over 38 sequential slices in descending order (slice thickness = 4 mm, slice gap = 1 mm), with the inferior-most slices prescribed at vertebral level T1 (TR = 2.5 s, TE = 30 ms, flip angle = 90°, ASSET factor = 2, FOV = 180 mm, acquisition matrix = 96×96, reconstruction matrix = 128×128). Static 0th, 1st & 2nd order shims were optimised. A spectral-spatial excitation pulse was used to excite only tissue water. Slice specific linear shims were implemented by adding 0.6 ms duration *x*-, *y*-, and *z*-gradient lobes after the excitation pulse. High-order shimming and *x, y*, and *z*-shimming were optimised over elliptical regions of interest (ROIs) covering the brain (for slices including the brain) or cord (for slices including the spinal cord). ROIs were drawn manually by the researcher (OSK or SM). To maintain consistency and avoid potential systematic differences in ROI drawing affecting test-retest estimates, the same researcher drew ROIs for both MRI sessions within participant.

Four dummy scans were acquired to enable the signal to reach steady-state, followed by 256 volumes. Full details of the acquisition sequence can be found in Tsivaka et al. (In prep). For 13 participants the manufacturer’s EPI internal reference option was used. The internal reference acquires four non-phase-encoded echoes before the EPI echo train, which are used to apply a phase correction to the EPI data. Upon further inspection of the data this was shown to contribute to slice misalignment (*y* direction) and thus the setting was disabled for the remaining participants. In order to keep the two MRI visits identical, however, the internal reference was used on both MRI visits for these 13 participants even after the issue was discovered.

### 2.4 Data preprocessing

Data were processed using Spinal Cord Toolbox (SCT) version 5.4 (De Leener et al., 2017), AFNI’s *3dWarpDrive* (Cox, 1996; Cox & Hyde, 1997), and FSL version 6.0.4 (Jenkinson et al., 2012; Smith et al., 2004). Visual quality assurance was performed on raw data and at each stage of processing. Five scans acquired with an early version of the functional sequence using the internal reference (see above) had several slices come out of alignment with the rest of the spinal cord due to a shift in the anterior-posterior (EPI phase-encoding) axis. A custom in-house Matlab (Mathworks Inc.) script was used to move the slices back into alignment with the rest of the cord. Briefly, for each slice, a 1D projection along the anterior/posterior direction was calculated for each time-point by summing the voxels in the left/right direction across the spinal cord. The anterior/posterior shift was determined by calculating the maximum of the cross correlation of the projection at each time-point with the first time-point. The shift was the applied to the image data in a block circular manner. Only shifts by an integer number of voxels were applied to avoid the need for an extra interpolation step. This step was performed prior to any other preprocessing.

For all functional data, brainstem structures were separated from cervical volumes at the level of the odontoid process. Subsequently, spinal cord functional data were motion-corrected for *x*- and *y*-translations using an in-house implementation of AFNI’s *3dWarpDrive* following the steps in the Neptune Toolbox (https://neptunetoolbox.com/). Motion-corrected data were smoothed with an in-plane 2D Gaussian kernel with full width at half maximum (FWHM) of 2 mm using a custom in-house script relying on tools from AFNI and FSL, and bandpass filtered (0.01-0.1 Hz) using *fslmaths* (part of FSL).

Warping parameters for spatial normalisation were determined by segmenting and registering the functional data to the Polytechnique Aix-Marseille University and Montreal Neurological Institute 50 (PAM50) template (De Leener et al., 2018), via an intermediary subject-specific T2-weighted 3D volume. Specifically, *sct_deepseg_sc* (Gros et al., 2019) was used to segment the cord from the cerebrospinal fluid (CSF) on motion-corrected functional data and on T2-weighted structural image (*sct_propseg* (De Leener et al., 2014) was used for one participant’s T2-weighted data where *sct_deepseg_sc* algorithm failed to detect the cord). Manual intervention was needed for accurate segmentation of functional data and was performed in FSLeyes (McCarthy, 2022). Warping parameters for registration of functional data to the PAM50 template were created by combining warp parameters from: (1) registering structural T2-weighted image to functional data utilising manually created disc labels on both images and (2) registering the segmented cord from T2-weighted image to the PAM50 T2-weigthed template via *sct_register_to_template* (De Leener et al., 2018). These warps were applied to functional data via *sct_register_multimodal* (De Leener et al., 2018). Inverse warp parameters obtained from these steps were used to transform PAM50 template cerebrospinal fluid and white matter masks to participant functional space which were used in the physiological denoising step described below.

The Physiological Noise Modelling (PNM) toolbox (Brooks et al., 2008) was used to generate 33 slice-specific regressors accounting for physiological noise based on cardiac and respiratory traces, and CSF signal. A bandpass filter (identical to that used on the functional data, 0.01-0.1Hz) was applied to nuisance regressors (those generated by the PNM and motion regressors obtained from motion correction as described above) to avoid reintroducing noise into the timeseries (Bright et al., 2017). Regression of physiological noise (cardiac and respiratory), cerebrospinal fluid and white matter signal, and motion parameters, along with pre-whitening using FILM were performed in FEAT. The smoothed and filtered data (i.e. the residuals from the previous step) were used for subsequent analyses.

### 2.5 Temporal signal-to-noise ratio (tSNR)

tSNR was calculated on minimally processed resting-state data to avoid artificially inflating the measure. The data had undergone motion correction only (as described above), to remove the timecourse variability associated with in-scan motion and enable creating subject-specific spinal cord masks (see detailed description of steps taken in generating cord masks above). tSNR maps were created by dividing the mean functional image by its standard deviation. Mean tSNR was extracted for the whole cervical cord (C1-C8) using subject-specific cord masks and for segmental levels C5-C8 using probabilistic segmental masks from the PAM50 atlas (De Leener et al., 2018) warped to subject-space (binarized and thresholded at 30% likelihood of belonging to that spinal level).

tSNR was extracted for all complete datasets (complete resting-state acquisition on both MRI sessions, i.e. 28 participants/56 resting-state acquisitions) that passed all other quality assurance steps (see Figure 1 for details). Since there are no established guidelines on cut-offs for inclusion based on data quality in spinal fMRI, we opted for a minimum tSNR of 20 to ensure reliability estimates were not affected by poor data quality. Consequently, five participants (i.e. 10 resting-state acquisitions) were excluded from all further analyses due to low mean tSNR across the whole cervical cord (<20) on at least one study session.

### 2.6 Assessment of resting-state networks

#### 2.6.1 Definition of seed regions

Seed regions were derived from the PAM50 atlas (De Leener et al., 2018) and corresponded to the four grey matter horns (ventral/dorsal and left/right) of 5^th^, 6^th^, 7^th^, and 8^th^ segmental levels. To obtain these masks we: 1) thresholded the mask of each horn (left/right, dorsal/ventral) at 50% likelihood of belonging to that grey matter horn and binarized it, 2) thresholded probabilistic segmental level (spinal levels C5-C8) masks at 30% to avoid overlap between segments, 3) multiplied each horn mask by each segmental level mask. This resulted in 16 individual masks for seed regions reflecting left/right and dorsal/ventral horns at segmental levels C5, C6, C7, and C8 (Figure 2).

**Figure 2.**
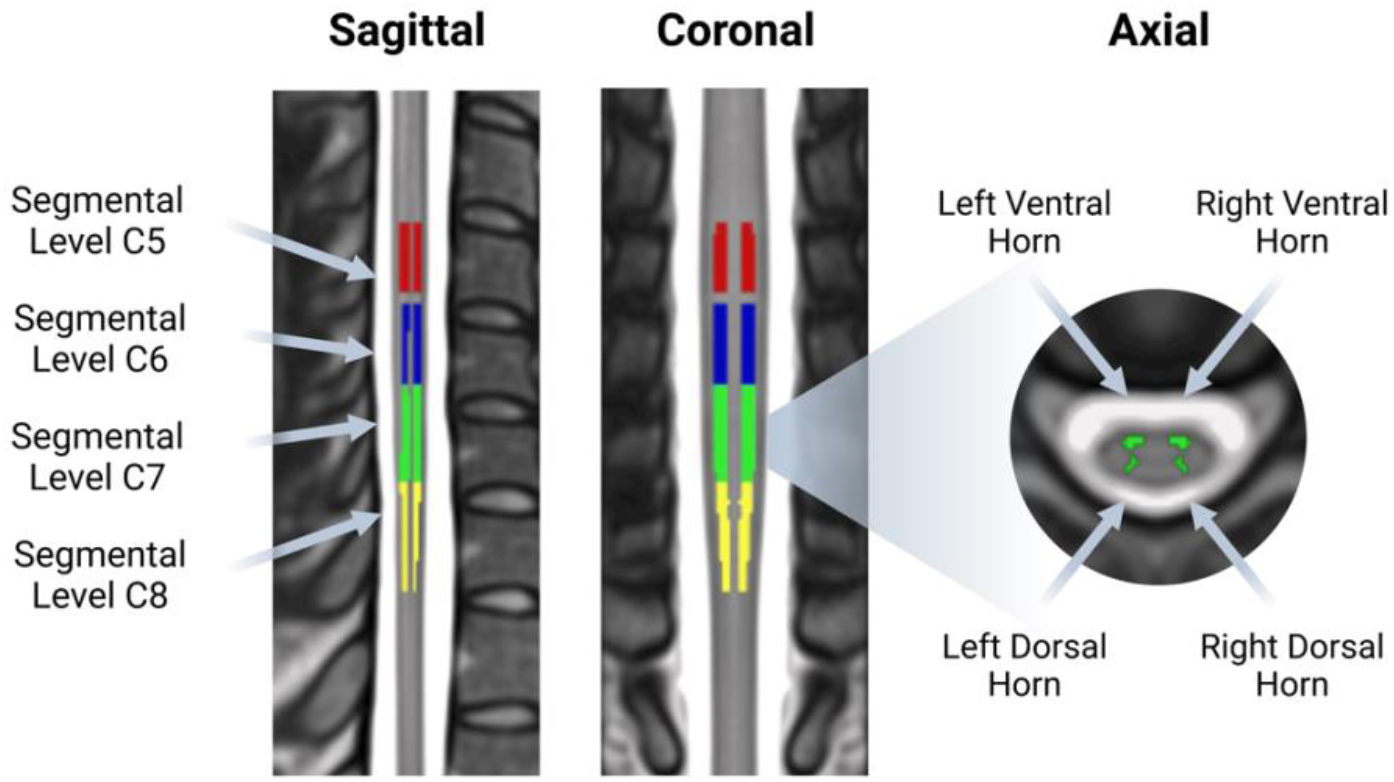
An illustration of the seed regions used in assessments of spinal cord resting-state networks. A total of 16 seeds were derived from the PAM50 atlas, corresponding to the four grey matter horns of the cord at spinal segmental levels C5 (red), C6 (blue), C7 (green), and C8 (yellow).

#### 2.6.2 Voxelwise connectivity

Mean timecourses extracted from these regions were used to estimate voxelwise functional connectivity maps within the cervical cord. For each subject, to assess both within- and between-segment connectivity all four seeds’ mean timecourses (left dorsal horn – L DH, right dorsal horn – R DH, left ventral horn – L VH, right ventral horn – R VH) for a given segmental level (C5, C6, C7, C8) were included in a single model estimated by FEAT. Consequently, a total of four models per subject, per session were run. COPE images from this stage were registered to PAM50 space using warp parameters generated during preprocessing (see above).

Spatial extent of resting-state networks at group level was assessed using *randomise* (Winkler et al., 2014) with threshold-free cluster enhancement (5000 permutations, *p* < 0.003 (*p* = 0.05, Bonferroni corrected for 16 individual seed regions)). This analysis was performed separately for each session.

#### 2.6.3 Seed-to-seed connectivity

In addition to the above preregistered voxelwise analysis, a more focused seed-to-seed correlation analysis was performed to assess the strength of connections between regions. Pearson correlations were computed between each pair of seed regions at subject-level using *numpy*.*corrcoef* function (Harris et al., 2020). The resultant correlation coefficients were *Z*-transformed using *numpy*.*arctanh* (Harris et al., 2020). Statistical significance at group-level was assessed using a one-sample *t*-test calculated using *scipy*.*stats*.*ttest_1samp* (Virtanen et al., 2020). A positive false discovery rate (FDR) was used to account for multiple comparisons (thresholded at *p* < 0.05, implemented with *statsmodels*.*stats*.*multitest*.*fdrcorrection* (Seabold & Perktold, 2010)). The analysis presented in the main text of the manuscript used data acquired on session 1 (see Supplementary Materials for corresponding analysis of data acquired on session 2).

### 2.7 Test-retest reliability

#### 2.7.1 Intraclass corelation coefficient (ICC)

To systematically evaluate the test-retest performance, inter-session intra-subject reliability was estimated using:

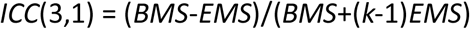

where *BMS* is the between-target mean squares, *EMS* is the error mean squares, and *k* is the number of repeated sessions.

ICC values were calculated for each voxel (i.e. voxelwise) using the locally-developed ICC toolbox (Caceres et al., 2009) running in Matlab version 9.5.0 (Mathworks Inc.). Intra-subject reliability was calculated for the whole cord and the complete activation network. The activation network was obtained using a one-sample *t*-test of the first session with a voxelwise *t*-statistic threshold of 3.5 (equivalent to *p* = 0.001) conducted in SPM (Caceres et al., 2009). ICC(3,1) was calculated for each COPE separately. Median ICC values are reported, defined as the reliability measure obtained from the median of the ICC distributions within regions. In addition to this pre-registered approach, additional ICC values were also computed to provide a more detailed understanding of the test-retest reliability of spinal resting-state data.

ICC(3,1) of the mean activation within a network was also computed. Mean signal was extracted from group-level maps obtained from *randomise* (as described above) using a binarized mask defined from activation map of session 1.

Additionally, ICC(3,1) values were calculated on the subject-level *Z*-scores describing each of the connections in the seed-to-seed analysis.

Finally, ICC(3,1) was calculated for tSNR values extracted from the whole cord and from segmental levels C5-C8 (see below). SPSS v28.0.1.1 with Python3 integration was used to calculate ICC values for mean activation within the network, seed-to-seed connectivities, and tSNR.

Following previous recommendations (Fleiss et al., 2013), ICC values will be categorised accordingly: <0.4 as poor, 0.4–0.59 as fair, 0.6–0.74 as good, and >0.75 as excellent. While a value of 1 indicates near-perfect agreement between the values of the test and retest sessions, a value of 0 would indicate that there was no agreement between the values of the test and retest sessions.

#### 2.7.2 Dice similarity coefficient (DSC)

Spatial consistency of spinal cord resting-state networks was evaluated using Dice similarity coefficient (DSC) (Dice, 1945) calculated using AFNI’s *3ddot* function. DSC was calculated separately for group- and subject-level maps. Mean DSC values for subject-level maps are reported.

DSC ranges from 0 to 1 with higher values indicating better overlap between two sets/maps. A value of 1 would thus correspond to perfect overlap, while a value of 0 would correspond to no overlap.

## 3 Results

### 3.1 tSNR

To assess signal quality, tSNR was extracted from minimally processed data (motion correction only) for all complete datasets (i.e. prior to excluding participants with mean tSNR across the whole cord < 20). Mean tSNR for the whole cord and segmental levels C5-C6 across sessions are shown in Table 1.

**Table 1.**
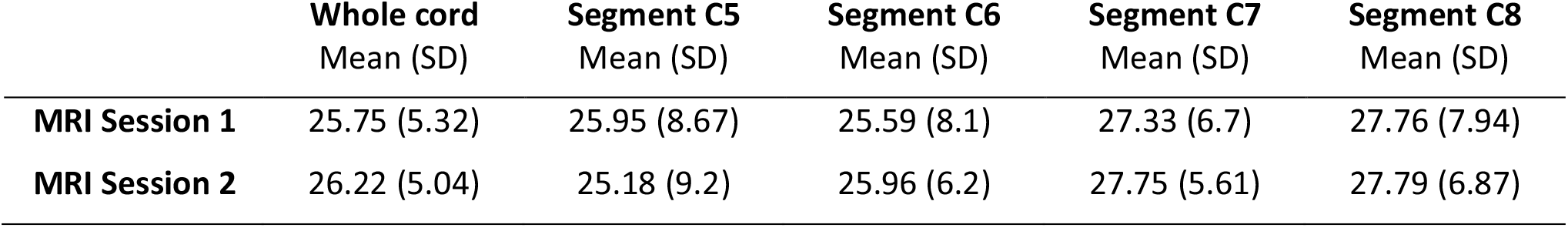
Temporal signal-to-noise ratio (tSNR) across whole cord and within spinal segmental levels C5-C8 for data acquired on MRI sessions 1 and 2. Data reported for N = 28, i.e. all complete datasets prior to excluding participants with tSNR < 20.

tSNR was stable across sessions both within the whole spinal cord (*t*(27) = -0.58, *p* = 0.568, *d* = 4.36, 95% CI [-0.48, 0.26]) and across segmental levels C5-C8 (*F*(1, 27) = 0, *p* = 0.989). Slightly higher tSNR was observed in lower segments (C7 and C8) than in higher segments (C5 and C6), however this difference was not statistically significant (*F*(1.93, 51.99) = 2.7, *p* = 0.078).

### 3.2 Assessment of resting-state networks

#### 3.2.1 Voxelwise connectivity

To assess the spatial extent of cervical spinal resting-state networks, we estimated voxelwise connectivity maps for each subject and session. This section describes the results of the analysis of data from session 1 (the corresponding analysis of session 2 data is provided in the Supplementary Materials). For each seed and segmental level, we observed a statistically significant organisation of spinal resting-state networks (*p* < 0.003). Each seed gave rise to a connectivity pattern that was largely confined to the segment, with sparser between-segment connections (Figure 3). While the spatial extent of clusters was similar across the four quadrants of each segment, we qualitatively observed a dorsal bias in functional connectivity of dorsal seeds and a ventral bias in functional connectivity of ventral seeds. Qualitatively, clusters estimated from session 2 data had highly similar spatial extent (see Supplementary Materials for results of session 2 data analysis and Supplementary Figure 2 for overlap between session 1 and session 2 maps).

**Figure 3.**
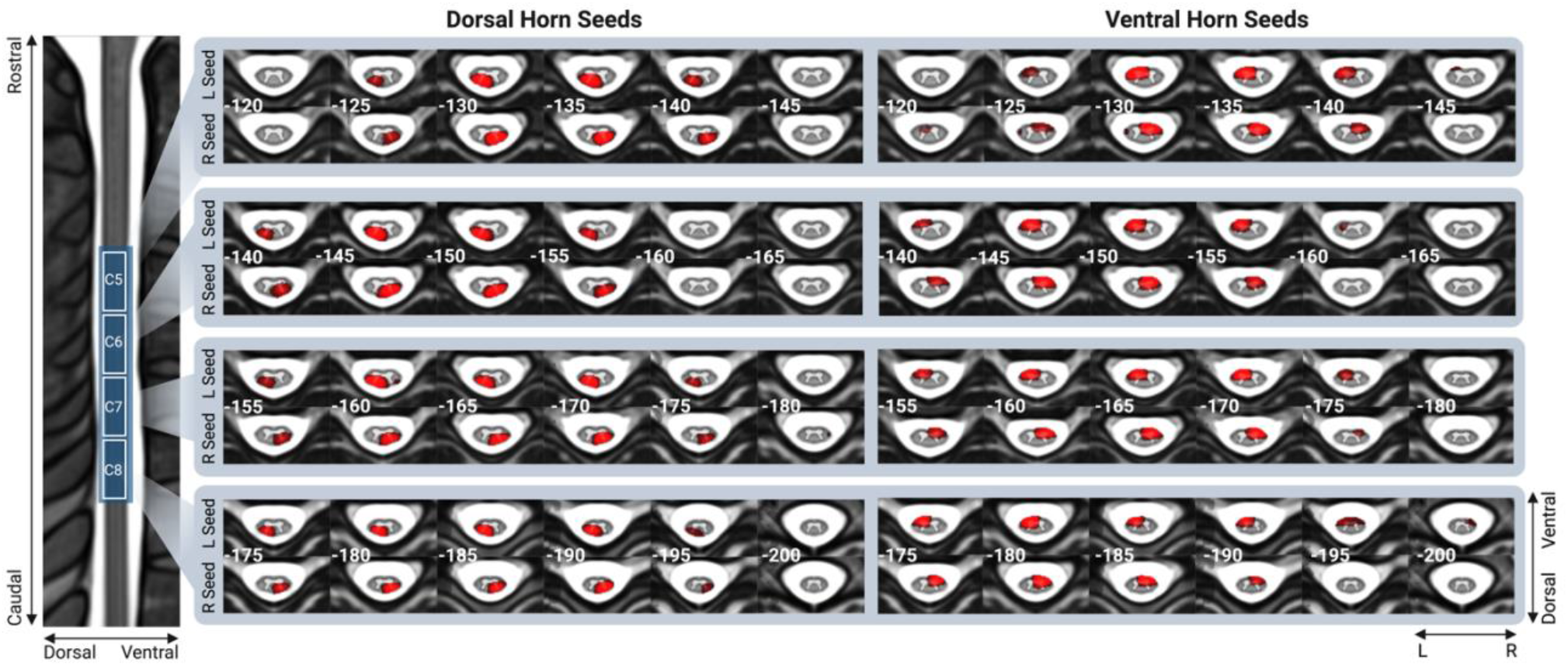
Resting-state networks obtained from voxelwise connectivity analysis for each of the four quadrants (ventral/dorsal and left/right) of segmental levels C5-C8 (data acquired on MRI session 1). Axial slices are marked with the *z* MNI coordinate. Each resting-state map was thresholded at *p*<0.003 (*p*=0.05, Bonferroni corrected for 16 individual seed regions).

#### 3.2.2 Seed-to-seed connectivity

To assess the strength of functional connections between horns of the cervical spinal cord, we conducted seed-to-seed correlations between each pair of seed regions on data acquired during session 1 (for results of the same analysis performed on session 2 data, see Supplementary Materials). A correlation matrix depicting cervical spinal cord connections is shown in Figure 4. On average, within segment, the strongest statistically significant positive correlations were observed within hemicord (i.e. left DH-VH and right DH-VH), followed by VH-VH and DH-DH connections, and DH-VH connections between hemicords (i.e. left DH – right VH, right DH – left VH). Weaker but statistically significant positive correlations were also observed between neighbouring segments, including DH-DH, VH-VH, as well as within and between hemicords. Finally, negative correlations were observed between the right VH of segment C8 and both left and right DH of segment C6. A similar pattern of results was observed in the analysis of data acquired during session 2 (see Supplementary Materials).

**Figure 4.**
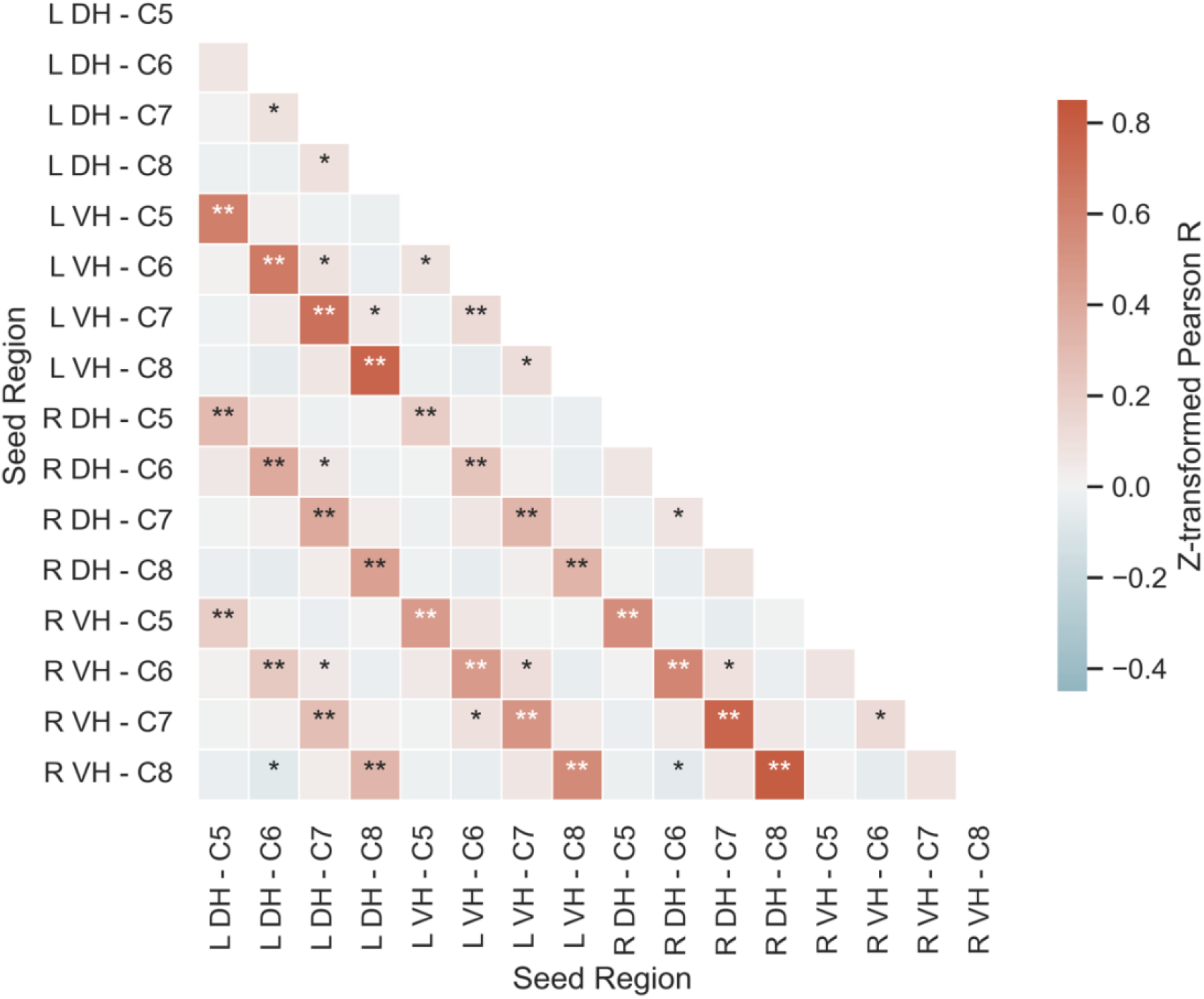
Seed-to-seed correlation matrix displaying z-transformed Person R. DH = Dorsal Horn, L = Left, VH = Ventral Horn, R = Right. *p < 0.05, **p < 0.001

### 3.3 Test-retest reliability

#### 3.3.1 ICC

ICC(3,1) was used to examine the test-retest reliability of cervical resting-state networks. ICC values for each resting-state network derived from voxelwise connectivity analysis are shown in Table 2 and for each of the seed-to-seed connectivities in Figure 5.

**Table 2.**
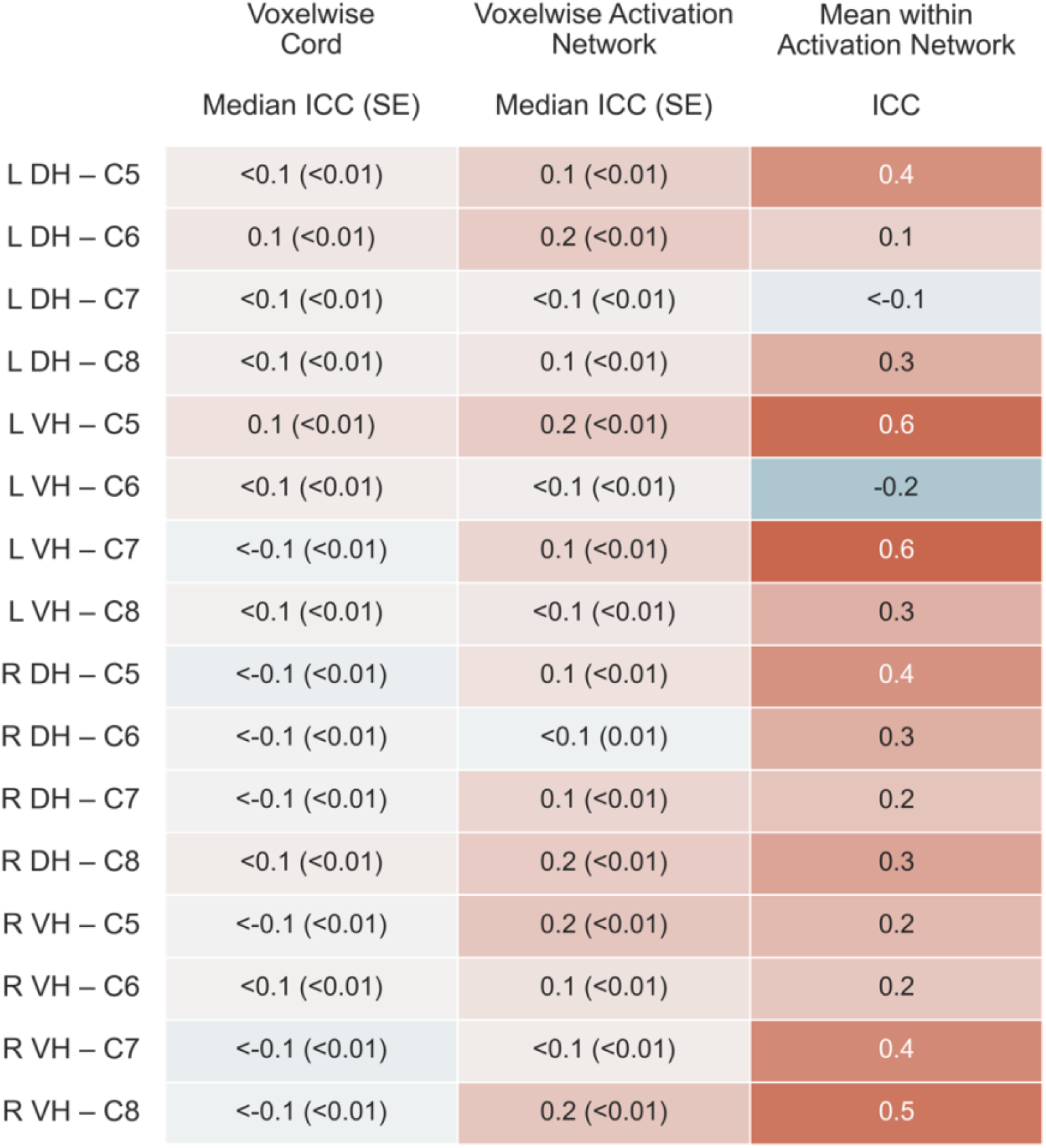
ICC(3,1) for each resting-state network. DH = Dorsal Horn, ICC = Intraclass Correlation Coefficient, L = Left, VH = Ventral Horn, R = Right.

**Figure 5.**
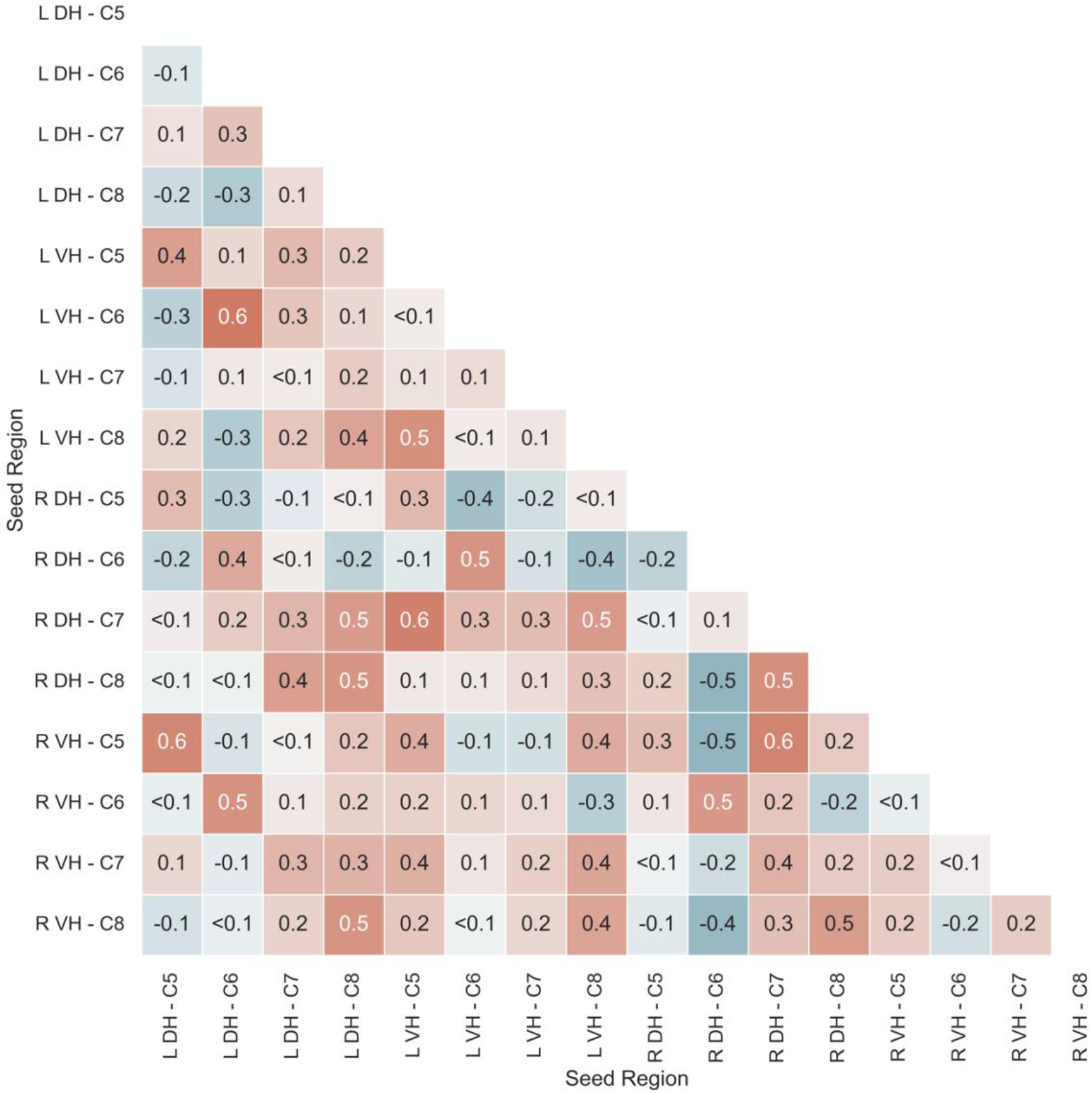
Matrix displaying ICC(3, 1) for each pair of seed regions. DH = Dorsal Horn, ICC = Intraclass Correlation Coefficient, L = Left, VH = Ventral Horn, R = Right.

On average voxelwise assessments of ICC in the entire cord (mean across networks ICC = <0.1 ± <0.1) and within the activation network defined based on MRI session 1 (mean across networks ICC = 0.1 ± <0.1) showed *poor* reliability across resting-state networks. ICCs for mean activation within each resting-state network showed better but still *poor* reliability (mean across networks ICC = 0.3 ± 0.2). Nonetheless, more variability in ICC values was observed, with some networks reaching *fair* (left and right DH networks at level C5 and right VH networks at levels C7 and C8) and *good* reliability (left VH networks at levels C5 and C6).

ICCs for connection strength across pairs of seed regions were variable. ICCs for a large portion of seed pairs (84%) were *poor*, however some reached *fair* (14%) and *good* (2%) levels. *Fair* and *good* ICCs were observed for connections both within and between spinal segmental levels and largely reflected either within (i.e. left DH-VH or right DH-VH) or between hemicord connectivity (i.e. left DH – right VH or right DH – left VH).

Finally, to assess the test-retest reliability of signal quality, ICC values were calculated for tSNR. Across the whole cervical spinal cord captured by our data, tSNR reliability was *good* (ICC = 0.7). Within segmental levels, tSNR reliability was *good* for segments C6 (ICC = 0.7), C7 (ICC = 0.6), and C8 (ICC = 0.7), and *fair* for segment C5 (ICC = 0.5).

#### 3.3.2 DSC

DSC assessed the spatial agreement of group- and subject-level resting-state maps between the two sessions. DSC for each network at group- and subject-level are shown in Table 3.

**Table 3.**
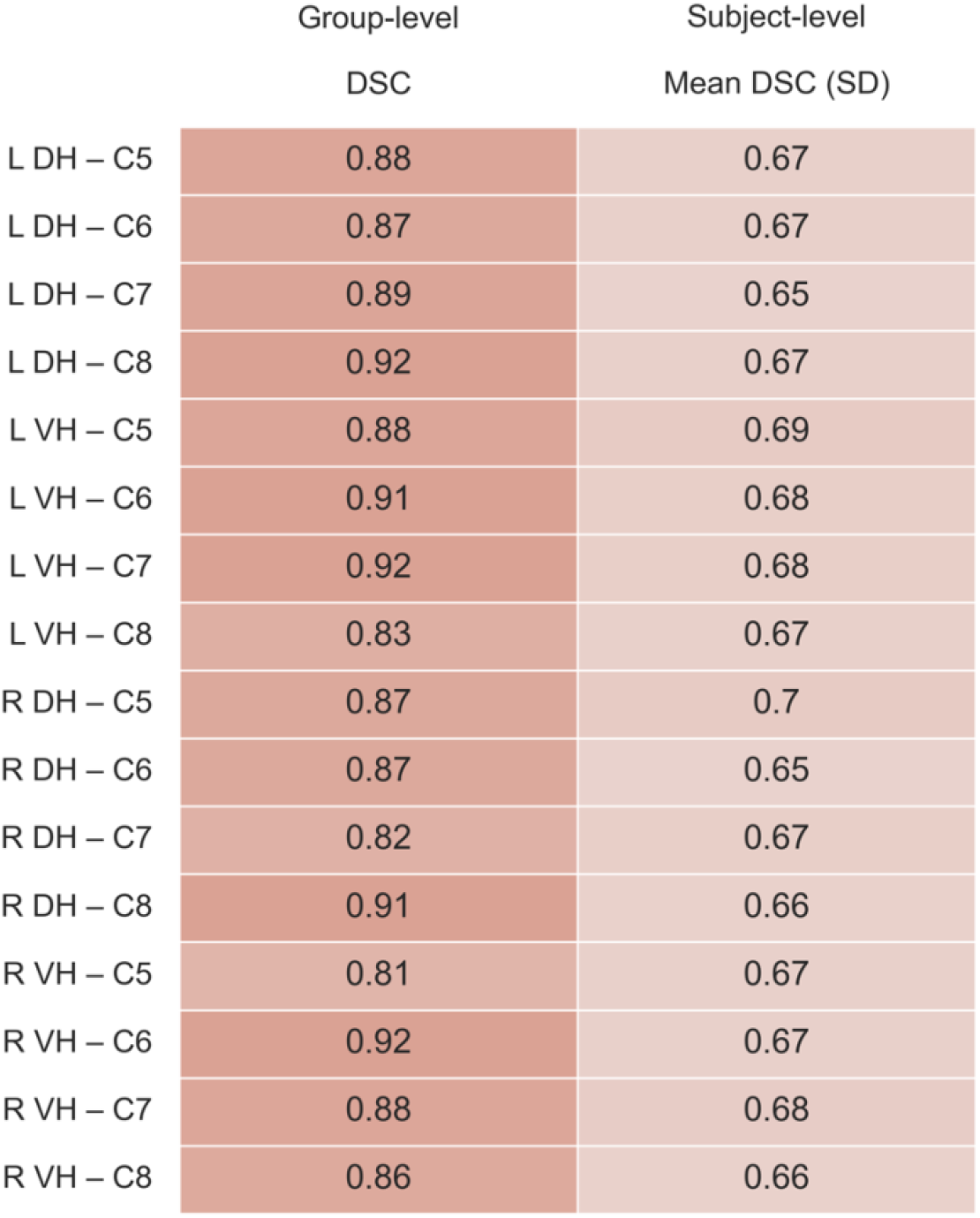
Group-level and mean subject-level DSC for each resting-state network. DH = Dorsal Horn, DSC = Dice Similarity Coefficient, L = Left, VH = Ventral Horn, R = Right.

Near-perfect agreement was observed in group-level maps (mean DSC = 0.88 ± 0.03) and good agreement was seen in subject-level maps (mean DSC = 0.67 ± 0.11).

## 4 Discussion

This study investigated cervical spinal cord resting-state networks and their test-retest reliability using a novel acquisition method. In mapping the spatial representation of resting-state networks, we observed distinct unilateral dorsal (sensory) and ventral (motor) organisation that was largely confined in the rostro-caudal extent to each spinal segmental level, with more sparse connections between segments. By investigating connection strength between the horns of the cervical spine, we observed that the strongest connectivity was present within the hemicord (i.e. ipsilateral dorsal-ventral), followed by ventro-ventral and dorso-dorsal connections, and finally dorsal-ventral connections between the hemicords. Similar but weaker connectivity was also observed between segmental levels. The results of test-retest reliability of these networks were mixed. Reliability was *poor* when assessed on a voxelwise level, with more promising but inconsistent indications of reliability when examining the average signal within networks and connection strength. However, assessments of the spatial overlap of resting-state network maps between sessions showed near-perfect agreement, suggesting that these networks are characterised by a consistent spatial representation over time.

The first aim of this study was to quantify the spatial extent of spinal cervical resting-state networks. Our findings of dorsal and ventral bias in the spatial representations of resting-state networks in the cervical spine are in line with our predictions and complement previous investigations characterising the intrinsic activity of the spinal cord (Barry et al., 2014, 2016; Eippert et al., 2017; Kong et al., 2014; Vahdat et al., 2020). In fact, the emergence of distinct sensory (dorsal) and motor (ventral) networks within the cervical spine has been demonstrated with several different analytical approaches, including data-driven independent component analysis (Kong et al., 2014; San Emeterio Nateras et al., 2016) and hypothesis-driven temporal correlation between regions of interest (Barry et al., 2014, 2016; Eippert et al., 2017). Further, these networks have been observed both at conventional MR field strength (3 T) (Eippert et al., 2017; Kong et al., 2014; Liu et al., 2016; San Emeterio Nateras et al., 2016; Vahdat et al., 2020) and at ultra-high field (7 T) (Barry et al., 2014, 2016). Here, we further confirm the presence of the previously reported dorso-dorsal and ventro-ventral cross-talk (Barry et al., 2014, 2016; Eippert et al., 2017) with seed-to-seed correlations and further show the emergence of unilateral dorsal and ventral networks (Kong et al., 2014) with voxelwise analyses. Our findings support the notion that these networks reflect intrinsic spinal activity, which mirrors the functional neuroanatomy of the spinal cord.

In addition to the distinct dorsal and ventral networks, we observed a strong within-hemicord (i.e. ipsilateral) connectivity between dorsal and ventral horns of the cervical spine. This is in contrast to previous reports of weak dorsal-ventral connectivity within the hemicord (Barry et al., 2014; Eippert et al., 2017). Nonetheless, strong within-hemicord connectivity between dorsal and ventral horns was observed in non-human primates (Chen et al., 2015) and in one study of a small groups of healthy adult volunteers (Weber et al., 2018). Furthermore, dorsal-ventral connectivity was also observed in some participants at ultra-high field, however, these results were not consistent and did not emerge at group level (Barry et al., 2014). Dorsal-ventral connectivity may represent a distinct sensory-motor spinal network, which could support motor reflexes and other more lateralised processing (Chen et al., 2015; Harrison et al., 2021). Indeed, anatomical spinal circuits that connect ipsilateral dorsal and ventral horns, including the monosynaptic stretch reflex and nociceptive withdrawal reflex, are well documented (Pierrot-Deseilligny & Burke, 2012). Nonetheless, given the close proximity of ipsilateral dorsal and ventral horns and the likely influence of fMRI acquisition parameters and data processing steps on the detectability of within-hemicord connectivity, further study is needed to establish whether these anatomical circuits contribute to a tertiary spinal resting-state network.

Similar to previous studies (Kinany et al., 2020; Kong et al., 2014; San Emeterio Nateras et al., 2016), we observed that spinal resting-state networks were largely limited in the rostro-caudal extent, mirroring the segmental organisation of the spinal cord. However, we also observed sparse between-segment connections. Intersegmental connectivity has been reported previously (Eippert et al., 2017; Harita & Stroman, 2017; Ioachim et al., 2019; San Emeterio Nateras et al., 2016; Vahdat et al., 2020) and is thought to reflect ascending sensory and descending motor pathways. In line with our findings, others have reported a decrease of connectivity beyond one vertebral level (Harita & Stroman, 2017; Liu et al., 2016; San Emeterio Nateras et al., 2016; Weber et al., 2018) and, in some cases, weak anti-correlation between regions of different segmental levels (Kinany et al., 2020; Kong et al., 2014). This pattern of results was also observed in this study, with an anti-correlation between right ventral horn at C8 and both ipsilateral and contralateral dorsal horn of segment C6. Such negative relationships may reflect processes related to intersegmental inhibition, perhaps contributing to reflexive actions, proprioception, and nociception (Friesen & Cang, 2001; McBain et al., 2016).

Our second aim was to assess whether cervical spinal resting-state networks could be reliably detected across different scanning sessions. The mixed findings observed in our reliability analysis are in contrast to our predictions and previous reports of *good* and *fair* reliability of resting-state connections in the cervical spine, albeit when tested within the same scanning session (Barry et al., 2016; Kaptan et al., 2022; Liu et al., 2016). Test-retest reliability is known to reduce with longer lag between sessions across various contexts (Calamia et al., 2013; Duff, 2012), including brain fMRI (Bennett & Miller, 2010, 2013) and specifically resting-state paradigms (Niu et al., 2020; Yang et al., 2022). Changes related to development, aging, learning, and attention, along with other neuroplastic processes likely underpin the biological reasons for poorer reliability in the long-term (Bennett & Miller, 2010, 2013). Furthermore, in cerebral fMRI, the highest reliability is usually achieved in data collected within the same scanning session (Shehzad et al., 2009; Wang et al., 2013), which likely reflects additional impact of scanner characteristics (An et al., 2017). Given that spinal cord fMRI acquisition is considerably more challenging than brain fMRI, with greater impact of baseline physiology and field inhomogeneities related to surrounding tissues, lower intersession test-retest estimates are to be expected.

In recent years, the reliability and reproducibility of neuroimaging results more broadly has been brought into question (Botvinik-Nezer et al., 2020; Poldrack et al., 2017), with largely mixed evidence of reliability across both task (Elliott et al., 2020; Kragel et al., 2021) and resting-state brain fMRI (Noble, Spann, et al., 2017; Noble et al., 2019). In fact, many estimates of brain resting-state connectivity achieve ICC values within the *poor* range (<0.4) across different resting-state metrics, including voxelwise and region-to-region connectivity (Noble et al., 2019; Noble, Scheinost, et al., 2017). Consequently, the test-retest estimates observed here for spinal cord resting-state networks are similar to those routinely observed in the brain. Furthermore, the spatial extents of these networks were similar across sessions. This suggests that while intensity changes in individual voxels and clusters may differ between sessions, the networks are characterised by a consistent spatial representation over time.

Aside from psychological influences, several factors have been identified, that contribute to low fMRI reliability, including poor tSNR (Bennett & Miller, 2010; Raemaekers et al., 2007), suboptimal data processing choices (Barry et al., 2016), and confounding effects of motion and/or other non-specific signal changes (Gorgolewski et al., 2013; Noble et al., 2019). The inherent challenges of acquiring spinal cord fMRI recordings, likely result in a compound effect of these factors, which may lead to somewhat lower test-retest reliability estimates than those of brain fMRI (Barry et al., 2016). The continued efforts to improve the quality of spinal cord recordings and finetune preprocessing pipelines will likely help to increase the reliability of spinal fMRI.

Nonetheless, it is important to recognise that high reliability does not always reflect data validity. For instance, it has been observed that correction for artefactual signal, such as motion and physiological noise, can lower test-retest reliability in the brain (Birn et al., 2014; Lipp et al., 2014; Noble et al., 2019; Noble, Spann, et al., 2017) and spinal cord (Kaptan et al., 2022). This likely represents more systematic properties of noise within the data (e.g. regular repetition of cardiac and/or respiratory processes, CSF pulsation leading to cord motion) compared to intrinsic activity within the cord, which may be characterised by more dynamic processes (Kinany et al., 2020). This is further supported by our observation of *good* reliability of the average tSNR of minimally processed data contrasting with lower reliability of resting-state networks estimated from the same data. Consequently, it is vital to consider data reliability and validity together and avoid data processing choices which, while boosting reliability, might have an undue effect on validity.

Most spinal-cord fMRI studies use *z*-shimming alone (Eippert et al., 2017; Kong et al., 2014; Vahdat et al., 2020). While not a primary intention of our study, we did observe that the *y*-shimming (and to a lesser extent *x*-shimming) gradients did provide additional signal recovery. One previous study has also reported dynamic *x*-, *y*-, and *z*-shimming (Islam et al., 2019), which differed from our implementation by applying the linear shimming gradients throughout the EPI acquisition for each slice rather than as gradient lobes. Additionally, we used spectral-spatial excitation pulses for our fMRI acquisition. Since these are designed to only excite water, no additional fat saturation pulses were required, which would have increased the TR needed to acquire images from 38 slices (or reduced the number of slices that could be acquired with the same TR). To date, spinal fMRI has been predominately implemented on Siemens scanners with only few exceptions (e.g. Islam et al., 2019). Our acquisition sequence uses a GE scanner platform and thus provides an alternative to the typically used Siemens-based methods.

The acquisition method described here achieved superior signal quality in comparison to reports describing other sequences used in the field to date, reaching an average tSNR of 26 across scanning sessions. This represents large gains over previously described methods, where average tSNR of spinal EPI data at 3 T typically ranges from 5 to 20 (Barry et al., 2018; Eippert et al., 2017; Kinany, Pirondini, Mattera, et al., 2022; Oliva et al., 2022; Powers et al., 2018). This boost in signal quality may be partly due to the slightly larger in-plane voxel size used in this study (1.4×1.4 mm compared to 1×1 mm typically used elsewhere (Eippert et al., 2017; Harita & Stroman, 2017; Kong et al., 2014; Liu et al., 2016; San Emeterio Nateras et al., 2016)). Aside from differences in voxel sizes, compared to brain fMRI, the low tSNR of spinal fMRI data is additionally driven by baseline physiology inducing spinal cord motion and CSF pulsation (Piché et al., 2009), and susceptibility artefacts arising from the distinct magnetic susceptibility profiles of surrounding tissues, resulting in signal dropout and image distortions (Saritas et al., 2014). While the tSNR achieved by our acquisition sequence remains lower than that of a typical brain EPI (tSNR of approximately 50 when calculated on minimally processed data) (Murphy et al., 2007; Oliva et al., 2022), it marks a step towards improving the quality of spinal fMRI recordings.

Several limitations are important to note in this study. Although we used a comparable voxel size (1.4×1.4 mm in-plane) to other spinal cord fMRI studies conducted at 3 T (Eippert et al., 2017; Harita & Stroman, 2017; Kong et al., 2014; Liu et al., 2016; San Emeterio Nateras et al., 2016), it needs to be noted that the small size of the spinal cord (approximately 10 mm in diameter with grey matter regions approximately 2-4 mm^2^ in-plane) (Harrison et al., 2021) calls for even finer spatial resolution in future studies. Although larger voxel size can improve signal-to-noise ratio, it can also lead to sampling signal from different structures within the same voxels. Similar issues arise from spatially smoothing the functional data. While smoothing increases tSNR and minimises variability in individual anatomy, it can lead to mixing of signal from distinct anatomical regions. This is particularly important to consider when investigating regions in close proximity (see above in relation to ipsilateral dorsal-ventral connectivity). Nonetheless, the correspondence of our findings and those of investigations conducted at higher field strength with smaller voxel size (0.31×0.31 mm in-plane) (Barry et al., 2014, 2016) and those not including spatial smoothing (Eippert et al., 2017; Kong et al., 2014), suggests that these were unlikely confounds in our data.

It is also important to consider that current best practices for spinal cord fMRI data modelling rely on assumptions that have been validated for cerebral fMRI but not studied in detail in the cord. For instance, early evidence suggests that frequencies higher than the conventional 0.08 Hz cut-off used for brain fMRI (Biswal et al., 1995), may be important drivers of spinal cord signalling (Barry et al., 2016). Here, we used bandpass filtering of 0.01-0.1Hz to allow for those higher frequencies, while keeping within the bounds of BOLD-validated frequency distribution. Nevertheless, the neurophysiological mechanisms underpinning assumptions crucial for fMRI data modelling, such as BOLD frequency distribution and haemodynamic response, require further study and validation in the cord.

Although we aimed to obtain 30 complete datasets, and indeed 37 participants completed one scanning session and 32 completed both sessions, the challenges associated with spinal cord fMRI acquisition and resultant data quality concerns meant that our final sample size was reduced to 23. Longer scanning time due to shimming optimisation, an additional anterior array coil resting on the participants neck and chest, head and neck positioning minimising neck curvature, and the use of external physiology monitoring equipment likely contributed to the discomfort associated with scanning, increased attrition rate, and led to higher in-scan motion. Further data exclusion was related to low tSNR and signal dropout, some of which may be a result of individual differences in the anatomy of surrounding tissues. High data attrition may be an inevitable attribute of spinal cord fMRI studies and needs to be accounted for during study design and recruitment.

Finally, our study investigated the test-retest reliability of cervical spinal resting-state networks across two separate sessions separated by several days or weeks, while previous studies looked at within-session reliability (Barry et al., 2016; Kaptan et al., 2022; Liu et al., 2016). However, a full characterisation of spinal cord fMRI reliability demands acquiring recordings from the same participants within the same session, as well as over days, weeks, months, and possibly years. Furthermore, combining recordings from the same subject across several sessions has been hypothesised to improve reliability alongside validity (Noble, Spann, et al., 2017). Such efforts in spinal cord fMRI may help to better understand the neurofunctional characteristics of spinal cord resting-state networks.

## 5 Conclusions

In this study, we demonstrate functional connectivity relationships in dorsal and ventral regions of the cervical cord using a novel acquisition method implemented on a GE platform. Importantly, our findings are in agreement with the known neuroanatomical and neurofunctional organisation of the spinal cord. Although the test-retest reliability of these networks was mixed, their spatial extent was highly reproducible across sessions, suggesting that these networks are characterised by a consistent spatial representation over time.

## Abbreviations

BOLD: Blood Oxygen Level Dependent
C: Cervical
DH: Dorsal Horn
DSC: Dice Similarity Coefficient
CSF: Cerebrospinal Fluid
EPI: Echo Planar Imaging
FDR: False Discovery Rate
fMRI: Functional Magnetic Resonance Imaging
FOV: Field of View
FWHM: Full Width at Half Maximum
GE: General Electric
ICC: Intraclass Correlation Coefficient
L: Left
NSAIDs: Non-Steroidal Anti-Inflammatory Drugs
PNM: Physiological Noise Modelling,
R: Right
ROI: Region of Interest
SCT: Spinal Cord Toolbox
SD: Standard Deviation
STAI: State Trait Anxiety Inventory
TE: Echo
TR: Repetition Time
tSNR: Temporal Signal-to-Noise Ratio
VH: Ventral Horn.

## Acknowledgements

We thank the radiographers for their help with MRI scanning and all volunteers for their participation in the study. We would also like to thank the University of Thessaly and Prof Ioannis Tsougos for supporting this project.

## CRediT author statement

Conceptualization: O.S.K., S.M., D.T., S.B.M., S.C.R.W., J.C.W.B., D.J.L., and M.A.H.

Data curation: O.S.K., S.M., and D.J.L.

Formal analysis: O.S.K., S.M., J.C.W.B., and M.A.H.

Funding acquisition: S.B.M., S.C.R.W., J.C.W.B., and M.A.H. Investigation: O.S.K., S.M., and M.A.H.

Methodology: O.S.K., S.M., D.T., J.C.W.B., D.J.L., and M.A.H.

Project administration: O.S.K., S.M., and M.A.H. Resources: O.S.K., S.M., and M.A.H.

Software: O.S.K., S.M., J.C.W.B., and D.J.L.

Supervision: M.A.H.

Validation: O.S.K., S.M., J.C.W.B., D.J.L., and M.A.H.

Visualization: O.S.K.

Writing - original draft: O.S.K. and D.J.L.

Writing - review & editing: O.S.K., S.M., D.T., S.C.R.W., J.C.W.B., D.J.L., and M.A.H.

## References

An, H. S., Moon, W.-J., Ryu, J.-K., Park, J. Y., Yun, W. S., Choi, J. W., Jahng, G.-H., & Park, J.-Y. (2017). Inter-vender and test-retest reliabilities of resting-state functional magnetic resonance imaging: Implications for multi-center imaging studies. Magnetic Resonance Imaging, 44, 125–130. https://doi.org/10.1016/j.mri.2017.09.001

Barry, R. L., Conrad, B. N., Smith, S. A., & Gore, J. C. (2018). A practical protocol for measurements of spinal cord functional connectivity. Scientific Reports, 8(1), Article 1. https://doi.org/10.1038/s41598-018-34841-6

Barry, R. L., Rogers, B. P., Conrad, B. N., Smith, S. A., & Gore, J. C. (2016). Reproducibility of resting state spinal cord networks in healthy volunteers at 7 Tesla. NeuroImage, 133, 31–40. https://doi.org/10.1016/j.neuroimage.2016.02.058

Barry, R. L., Smith, S. A., Dula, A. N., & Gore, J. C. (2014). Resting state functional connectivity in the human spinal cord. ELife, 3, e02812. https://doi.org/10.7554/eLife.02812

Bennett, C. M., & Miller, M. B. (2010). How reliable are the results from functional Magnetic Resonance Imaging? Annals of the New York Academy of Sciences, 1191, 133–155. https://doi.org/10.1111/j.1749-6632.2010.05446.x

Bennett, C. M., & Miller, M. B. (2013). fMRI reliability: Influences of task and experimental design. Cognitive, Affective, & Behavioral Neuroscience, 13(4), 690–702. https://doi.org/10.3758/s13415-013-0195-1

Bernstein, M. A., King, K. F., & Zhou, X. J. (2004). Basic Pulse Sequences. In M. A. Bernstein, K. F. King, & X. J. Zhou (Eds.), Handbook of MRI Pulse Sequences (pp. 579–647). Academic Press. https://doi.org/10.1016/B978-012092861-3/50021-2

Birn, R. M., Cornejo, M. D., Molloy, E. K., Patriat, R., Meier, T. B., Kirk, G. R., Nair, V. A., Meyerand, M. E., & Prabhakaran, V. (2014). The influence of physiological noise correction on test-retest reliability of resting-state functional connectivity. Brain Connectivity, 4(7), 511–522. https://doi.org/10.1089/brain.2014.0284

Biswal, B., Yetkin, F. Z., Haughton, V. M., & Hyde, J. S. (1995). Functional connectivity in the motor cortex of resting human brain using echo-planar MRI. Magnetic Resonance in Medicine, 34(4), 537–541. https://doi.org/10.1002/mrm.1910340409

Botvinik-Nezer, R., Holzmeister, F., Camerer, C. F., Dreber, A., Huber, J., Johannesson, M., Kirchler, M., Iwanir, R., Mumford, J. A., Adcock, R. A., Avesani, P., Baczkowski, B. M., Bajracharya, A., Bakst, L., Ball, S., Barilari, M., Bault, N., Beaton, D., Beitner, J., … Schonberg, T. (2020). Variability in the analysis of a single neuroimaging dataset by many teams. Nature, 1–7. https://doi.org/10.1038/s41586-020-2314-9

Bright, M. G., Tench, C. R., & Murphy, K. (2017). Potential pitfalls when denoising resting state fMRI data using nuisance regression. NeuroImage, 154, 159–168. https://doi.org/10.1016/j.neuroimage.2016.12.027

Brooks, J. C. W., Beckmann, C. F., Miller, K. L., Wise, R. G., Porro, C. A., Tracey, I., & Jenkinson, M. (2008). Physiological noise modelling for spinal functional magnetic resonance imaging studies. NeuroImage, 39(2), 680–692. https://doi.org/10.1016/j.neuroimage.2007.09.018

Caceres, A., Hall, D. L., Zelaya, F. O., Williams, S. C. R., & Mehta, M. A. (2009). Measuring fMRI reliability with the intra-class correlation coefficient. NeuroImage, 45(3), 758–768. https://doi.org/10.1016/j.neuroimage.2008.12.035

Calamia, M., Markon, K., & Tranel, D. (2013). The Robust Reliability of Neuropsychological Measures: Meta-Analyses of Test–Retest Correlations. The Clinical Neuropsychologist, 27(7), 1077–1105. https://doi.org/10.1080/13854046.2013.809795

Chen, L. M., Mishra, A., Yang, P.-F., Wang, F., & Gore, J. C. (2015). Injury alters intrinsic functional connectivity within the primate spinal cord. Proceedings of the National Academy of Sciences of the United States of America, 112(19), 5991–5996. https://doi.org/10.1073/pnas.1424106112

Cohen-Adad, J., Alonso-Ortiz, E., Abramovic, M., Arneitz, C., Atcheson, N., Barlow, L., Barry, R. L., Barth, M., Battiston, M., Büchel, C., Budde, M., Callot, V., Combes, A. J. E., De Leener, B., Descoteaux, M., de Sousa, P. L., Dostál, M., Doyon, J., Dvorak, A., … Xu, J. (2021). Generic acquisition protocol for quantitative MRI of the spinal cord. Nature Protocols, 16(10), 4611–4632. https://doi.org/10.1038/s41596-021-00588-0

Cox, R. W. (1996). AFNI: Software for analysis and visualization of functional magnetic resonance neuroimages. Computers and Biomedical Research, an International Journal, 29(3), 162–173. https://doi.org/10.1006/cbmr.1996.0014

Cox, R. W., & Hyde, J. S. (1997). Software tools for analysis and visualization of fMRI data. NMR in Biomedicine, 10(4–5), 171–178. https://doi.org/10.1002/(sici)1099-1492(199706/08)10:4/5<171::aid-nbm453>3.0.co;2-l

De Leener, B., Fonov, V. S., Collins, D. L., Callot, V., Stikov, N., & Cohen-Adad, J. (2018). PAM50: Unbiased multimodal template of the brainstem and spinal cord aligned with the ICBM152 space. NeuroImage, 165, 170–179. https://doi.org/10.1016/j.neuroimage.2017.10.041

De Leener, B., Kadoury, S., & Cohen-Adad, J. (2014). Robust, accurate and fast automatic segmentation of the spinal cord. NeuroImage, 98, 528–536. https://doi.org/10.1016/j.neuroimage.2014.04.051

De Leener, B., Lévy, S., Dupont, S. M., Fonov, V. S., Stikov, N., Louis Collins, D., Callot, V., & Cohen-Adad, J. (2017). SCT: Spinal Cord Toolbox, an open-source software for processing spinal cord MRI data. NeuroImage, 145, 24–43. https://doi.org/10.1016/j.neuroimage.2016.10.009

Dice, L. R. (1945). Measures of the Amount of Ecologic Association Between Species. Ecology, 26(3), 297–302. https://doi.org/10.2307/1932409

Drysdale, A. T., Grosenick, L., Downar, J., Dunlop, K., Mansouri, F., Meng, Y., Fetcho, R. N., Zebley, B., Oathes, D. J., Etkin, A., Schatzberg, A. F., Sudheimer, K., Keller, J., Mayberg, H. S., Gunning, F. M., Alexopoulos, G. S., Fox, M. D., Pascual-Leone, A., Voss, H. U., … Liston, C. (2017). Resting-state connectivity biomarkers define neurophysiological subtypes of depression. Nature Medicine, 23(1), Article 1. https://doi.org/10.1038/nm.4246

Duff, K. (2012). Evidence-based indicators of neuropsychological change in the individual patient: Relevant concepts and methods. Archives of Clinical Neuropsychology: The Official Journal of the National Academy of Neuropsychologists, 27(3), 248–261. https://doi.org/10.1093/arclin/acr120

Eippert, F., Kong, Y., Winkler, A. M., Andersson, J. L., Finsterbusch, J., Büchel, C., Brooks, J. C. W., & Tracey, I. (2017). Investigating resting-state functional connectivity in the cervical spinal cord at 3 T. Neuroimage, 147, 589–601. https://doi.org/10.1016/j.neuroimage.2016.12.072

Elliott, M. L., Knodt, A. R., Ireland, D., Morris, M. L., Poulton, R., Ramrakha, S., Sison, M. L., Moffitt, T. E., Caspi, A., & Hariri, A. R. (2020). What Is the Test-Retest Reliability of Common Task-Functional MRI Measures? New Empirical Evidence and a Meta-Analysis. Psychological Science, 31(7), 792–806. https://doi.org/10.1177/0956797620916786

Finsterbusch, J., Eippert, F., & Büchel, C. (2012). Single, slice-specific z-shim gradient pulses improve T2*-weighted imaging of the spinal cord. NeuroImage, 59(3), 2307–2315. https://doi.org/10.1016/j.neuroimage.2011.09.038

Fleiss, J. L., Levin, B., & Paik, M. C. (2013). Statistical methods for rates and proportions. John Wiley & Sons.

Friesen, W. O., & Cang, J. (2001). Sensory and central mechanisms control intersegmental coordination. Current Opinion in Neurobiology, 11(6), 678–683. https://doi.org/10.1016/S0959-4388(01)00268-9

Gorgolewski, K. J., Storkey, A. J., Bastin, M. E., Whittle, I., & Pernet, C. (2013). Single subject fMRI test–retest reliability metrics and confounding factors. NeuroImage, 69, 231–243. https://doi.org/10.1016/j.neuroimage.2012.10.085

Gros, C., De Leener, B., Badji, A., Maranzano, J., Eden, D., Dupont, S. M., Talbott, J., Zhuoquiong, R., Liu, Y., Granberg, T., Ouellette, R., Tachibana, Y., Hori, M., Kamiya, K., Chougar, L., Stawiarz, L., Hillert, J., Bannier, E., Kerbrat, A., … Cohen-Adad, J. (2019). Automatic segmentation of the spinal cord and intramedullary multiple sclerosis lesions with convolutional neural networks. NeuroImage, 184, 901–915. https://doi.org/10.1016/j.neuroimage.2018.09.081

Harita, S., & Stroman, P. W. (2017). Confirmation of resting-state BOLD fluctuations in the human brainstem and spinal cord after identification and removal of physiological noise. Magnetic Resonance in Medicine, 78(6), 2149–2156. https://doi.org/10.1002/mrm.26606

Harris, C. R., Millman, K. J., van der Walt, S. J., Gommers, R., Virtanen, P., Cournapeau, D., Wieser, E., Taylor, J., Berg, S., Smith, N. J., Kern, R., Picus, M., Hoyer, S., van Kerkwijk, M. H., Brett, M., Haldane, A., del Río, J. F., Wiebe, M., Peterson, P., … Oliphant, T. E. (2020). Array programming with NumPy. Nature, 585(7825), Article 7825. https://doi.org/10.1038/s41586-020-2649-2

Harrison, O. K., Guell, X., Klein-Flügge, M. C., & Barry, R. L. (2021). Structural and resting state functional connectivity beyond the cortex. NeuroImage, 240, 118379. https://doi.org/10.1016/j.neuroimage.2021.118379

Ioachim, G., Powers, J. M., & Stroman, P. W. (2019). Comparing Coordinated Networks Across the Brainstem and Spinal Cord in the Resting State and Altered Cognitive State. Brain Connectivity, 9(5), 415–424. https://doi.org/10.1089/brain.2018.0659

Islam, H., Law, C. S. W., Weber, K. A., Mackey, S. C., & Glover, G. H. (2019). Dynamic per slice shimming for simultaneous brain and spinal cord fMRI. Magnetic Resonance in Medicine, 81(2), 825–838. https://doi.org/10.1002/mrm.27388

Jenkinson, M., Beckmann, C. F., Behrens, T. E. J., Woolrich, M. W., & Smith, S. M. (2012). FSL. NeuroImage, 62(2), 782–790. https://doi.org/10.1016/j.neuroimage.2011.09.015

Kaptan, M., Horn, U., Vannesjo, S. J., Mildner, T., Weiskopf, N., Finsterbusch, J., Brooks, J. C. W., & Eippert, F. (2022). Reliability of resting-state functional connectivity in the human spinal cord: Assessing the impact of distinct noise sources (p. 2022.12.23.521768). bioRxiv. https://doi.org/10.1101/2022.12.23.521768

Kinany, N., Pirondini, E., Mattera, L., Martuzzi, R., Micera, S., & Van De Ville, D. (2022). Towards reliable spinal cord fMRI: Assessment of common imaging protocols. NeuroImage, 250, 118964. https://doi.org/10.1016/j.neuroimage.2022.118964

Kinany, N., Pirondini, E., Micera, S., & Van De Ville, D. (2020). Dynamic Functional Connectivity of Resting-State Spinal Cord fMRI Reveals Fine-Grained Intrinsic Architecture. Neuron, 108(3), 424–435.e4. https://doi.org/10.1016/j.neuron.2020.07.024

Kinany, N., Pirondini, E., Micera, S., & Van De Ville, D. (2022). Spinal Cord fMRI: A New Window into the Central Nervous System. The Neuroscientist, 10738584221101828. https://doi.org/10.1177/10738584221101827

Kong, Y., Eippert, F., Beckmann, C. F., Andersson, J., Finsterbusch, J., Büchel, C., Tracey, I., & Brooks, J. C. W. (2014). Intrinsically organized resting state networks in the human spinal cord. Proceedings of the National Academy of Sciences, 111(50), 18067–18072. https://doi.org/10.1073/pnas.1414293111

Kowalczyk, O. S., Medina, S., Howard, M. A., Tsivaka, D., Lythgoe, D. J., & Brooks, J. C. W. (2021). Examining test-retest reliability of evoked response and resting-state functional MRI endpoints in human brain and cervical spine. https://doi.org/10.17605/osf.io/u8rbq

Kragel, P. A., Han, X., Kraynak, T. E., Gianaros, P. J., & Wager, T. D. (2021). Functional MRI Can Be Highly Reliable, but It Depends on What You Measure: A Commentary on Elliott et al. (2020). Psychological Science, 32(4), 622–626. https://doi.org/10.1177/0956797621989730

Lipp, I., Murphy, K., Wise, R. G., & Caseras, X. (2014). Understanding the contribution of neural and physiological signal variation to the low repeatability of emotion-induced BOLD responses. NeuroImage, 86, 335–342. https://doi.org/10.1016/j.neuroimage.2013.10.015

Liu, X., Zhou, F., Li, X., Qian, W., Cui, J., Zhou, I. Y., Luk, K. D. K., Wu, Ed. X., & Hu, Y. (2016). Organization of the intrinsic functional network in the cervical spinal cord: A resting state functional MRI study. Neuroscience, 336, 30–38. https://doi.org/10.1016/j.neuroscience.2016.08.042

McBain, R. A., Taylor, J. L., Gorman, R. B., Gandevia, S. C., & Butler, J. E. (2016). Human intersegmental reflexes from intercostal afferents to scalene muscles. Experimental Physiology, 101(10), 1301–1308. https://doi.org/10.1113/EP085907

McCarthy, P. (2022). FSLeyes (1.5.0). Zenodo. https://doi.org/10.5281/zenodo.7038115

Murphy, K., Bodurka, J., & Bandettini, P. A. (2007). How long to scan? The relationship between fMRI temporal signal to noise and necessary scan duration. NeuroImage, 34(2), 565–574. https://doi.org/10.1016/j.neuroimage.2006.09.032

Niu, Y., Sun, J., Wang, B., Hussain, W., Fan, C., Cao, R., Zhou, M., & Xiang, J. (2020). Comparing Test-Retest Reliability of Entropy Methods: Complexity Analysis of Resting-State fMRI. IEEE Access, 8, 124437–124450. https://doi.org/10.1109/ACCESS.2020.3005906

Noble, S., Scheinost, D., & Constable, R. T. (2019). A decade of test-retest reliability of functional connectivity: A systematic review and meta-analysis. NeuroImage, 203, 116157. https://doi.org/10.1016/j.neuroimage.2019.116157

Noble, S., Scheinost, D., Finn, E. S., Shen, X., Papademetris, X., McEwen, S. C., Bearden, C. E., Addington, J., Goodyear, B., Cadenhead, K. S., Mirzakhanian, H., Cornblatt, B. A., Olvet, D. M., Mathalon, D. H., McGlashan, T. H., Perkins, D. O., Belger, A., Seidman, L. J., Thermenos, H., … Constable, R. T. (2017). Multisite reliability of MR-based functional connectivity. NeuroImage, 146, 959–970. https://doi.org/10.1016/j.neuroimage.2016.10.020

Noble, S., Spann, M. N., Tokoglu, F., Shen, X., Constable, R. T., & Scheinost, D. (2017). Influences on the Test–Retest Reliability of Functional Connectivity MRI and its Relationship with Behavioral Utility. Cerebral Cortex, 27(11), 5415–5429. https://doi.org/10.1093/cercor/bhx230

Oldfield, R. C. (1971). The assessment and analysis of handedness: The Edinburgh inventory. Neuropsychologia, 9(1), 97–113. https://doi.org/10.1016/0028-3932(71)90067-4

Oliva, V., Hartley-Davies, R., Moran, R., Pickering, A. E., & Brooks, J. C. (2022). Simultaneous brain, brainstem, and spinal cord pharmacological-fMRI reveals involvement of an endogenous opioid network in attentional analgesia. ELife, 11, e71877. https://doi.org/10.7554/eLife.71877

Pfannmöller, J., & Lotze, M. (2019). Review on biomarkers in the resting-state networks of chronic pain patients. Brain and Cognition, 131, 4–9. https://doi.org/10.1016/j.bandc.2018.06.005

Piché, M., Cohen-Adad, J., Nejad, M. K., Perlbarg, V., Xie, G., Beaudoin, G., Benali, H., & Rainville, P. (2009). Characterization of cardiac-related noise in fMRI of the cervical spinal cord. Magnetic Resonance Imaging, 27(3), 300–310. https://doi.org/10.1016/j.mri.2008.07.019

Pierrot-Deseilligny, E., & Burke, D. (2012). The Circuitry of the Human Spinal Cord: Spinal and Corticospinal Mechanisms of Movement. Cambridge University Press.

Poldrack, R. A., Baker, C. I., Durnez, J., Gorgolewski, K. J., Matthews, P. M., Munafò, M. R., Nichols, T. E., Poline, J.-B., Vul, E., & Yarkoni, T. (2017). Scanning the horizon: Towards transparent and reproducible neuroimaging research. Nature Reviews Neuroscience, 18(2), 115–126.

Powers, J. M., Ioachim, G., & Stroman, P. W. (2018). Ten Key Insights into the Use of Spinal Cord fMRI. Brain Sciences, 8(9), E173. https://doi.org/10.3390/brainsci8090173

Raemaekers, M., Vink, M., Zandbelt, B., van Wezel, R. J. A., Kahn, R. S., & Ramsey, N. F. (2007). Test-retest reliability of fMRI activation during prosaccades and antisaccades. NeuroImage, 36(3), 532–542. https://doi.org/10.1016/j.neuroimage.2007.03.061

San Emeterio Nateras, O., Yu, F., Muir, E. R., Bazan, C., Franklin, C. G., Li, W., Li, J., Lancaster, J. L., & Duong, T. Q. (2016). Intrinsic Resting-State Functional Connectivity in the Human Spinal Cord at 3.0 T. Radiology, 279(1), 262–268. https://doi.org/10.1148/radiol.2015150768

Saritas, E. U., Holdsworth, S. J., & Bammer, R. (2014). Chapter 2.3—Susceptibility Artifacts. In J. Cohen-Adad & C. A. M. Wheeler-Kingshott (Eds.), Quantitative MRI of the Spinal Cord (pp. 91–105). Academic Press. https://doi.org/10.1016/B978-0-12-396973-6.00007-1

Seabold, S., & Perktold, J. (2010). Statsmodels: Econometric and Statistical Modeling with Python. 92–96. https://doi.org/10.25080/Majora-92bf1922-011

Shehzad, Z., Kelly, A. M. C., Reiss, P. T., Gee, D. G., Gotimer, K., Uddin, L. Q., Lee, S. H., Margulies, D. S., Roy, A. K., Biswal, B. B., Petkova, E., Castellanos, F. X., & Milham, M. P. (2009). The Resting Brain: Unconstrained yet Reliable. Cerebral Cortex (New York, NY), 19(10), 2209–2229. https://doi.org/10.1093/cercor/bhn256

Smith, S. M., Jenkinson, M., Woolrich, M. W., Beckmann, C. F., Behrens, T. E. J., Johansen-Berg, H., Bannister, P. R., De Luca, M., Drobnjak, I., Flitney, D. E., Niazy, R. K., Saunders, J., Vickers, J., Zhang, Y., De Stefano, N., Brady, J. M., & Matthews, P. M. (2004). Advances in functional and structural MR image analysis and implementation as FSL. NeuroImage, 23 Suppl 1, S208–219. https://doi.org/10.1016/j.neuroimage.2004.07.051

Spielberger, C. D., Gonzalez-Reigosa, F., Martinez-Urrutia, A., Natalicio, L. F., & Natalicio, D. S. (1971). The State-Trait Anxiety Inventory. Revista Interamericana De Psicología/Interamerican Journal of Psychology, 5(3–4). https://journal.sipsych.org/index.php/IJP/article/view/620

Taylor, J. J., Kurt, H. G., & Anand, A. (2021). Resting State Functional Connectivity Biomarkers of Treatment Response in Mood Disorders: A Review. Frontiers in Psychiatry, 12. https://www.frontiersin.org/articles/10.3389/fpsyt.2021.565136

Tinnermann, A., Büchel, C., & Cohen-Adad, J. (2021). Cortico-spinal imaging to study pain. NeuroImage, 224, 117439. https://doi.org/10.1016/j.neuroimage.2020.117439

Vahdat, S., Khatibi, A., Lungu, O., Finsterbusch, J., Büchel, C., Cohen-Adad, J., Marchand-Pauvert, V., & Doyon, J. (2020). Resting-state brain and spinal cord networks in humans are functionally integrated. PLoS Biology, 18(7), e3000789. https://doi.org/10.1371/journal.pbio.3000789

Virtanen, P., Gommers, R., Oliphant, T. E., Haberland, M., Reddy, T., Cournapeau, D., Burovski, E., Peterson, P., Weckesser, W., Bright, J., van der Walt, S. J., Brett, M., Wilson, J., Millman, K. J., Mayorov, N., Nelson, A. R. J., Jones, E., Kern, R., Larson, E., … van Mulbregt, P. (2020). SciPy 1.0: Fundamental algorithms for scientific computing in Python. Nature Methods, 17(3), Article 3. https://doi.org/10.1038/s41592-019-0686-2

Wang, X., Jiao, Y., Tang, T., Wang, H., & Lu, Z. (2013). Investigating univariate temporal patterns for intrinsic connectivity networks based on complexity and low-frequency oscillation: A test–retest reliability study. Neuroscience, 254, 404–426. https://doi.org/10.1016/j.neuroscience.2013.09.009

Weber, K. A., Sentis, A. I., Bernadel-Huey, O. N., Chen, Y., Wang, X., Parrish, T. B., & Mackey, S. (2018). Thermal Stimulation Alters Cervical Spinal Cord Functional Connectivity in Humans. Neuroscience, 369, 40–50. https://doi.org/10.1016/j.neuroscience.2017.10.035

Winkler, A. M., Ridgway, G. R., Webster, M. A., Smith, S. M., & Nichols, T. E. (2014). Permutation inference for the general linear model. NeuroImage, 92, 381–397. https://doi.org/10.1016/j.neuroimage.2014.01.060

Yang, L., Wei, J., Li, Y., Wang, B., Guo, H., Yang, Y., & Xiang, J. (2022). Test–Retest Reliability of Synchrony and Metastability in Resting State fMRI. Brain Sciences, 12(1), Article 1. https://doi.org/10.3390/brainsci12010066

