## Supplementary Materials for "A novel implementation of spinal fMRI demonstrates segmental organisation of functionally connected networks in the cervical spinal cord: A test-retest reliability study"

### 1. Voxelwise connectivity analysis for data acquired during MRI session 2

#### 1.1 Methods

Seed-based voxelwise connectivity analysis of data acquired during MRI session 2 followed the same steps as that of MRI session 1 (described in the main text). Mean timecourses extracted from regions of interest (L/R DH/VH of each segmental level from C5 to C8) were used to provide voxelwise estimates of functional connectivity within the entire cervical cord. Subject-level modelling was performed in FEAT by including all four seeds’ timecourses (L DH, R DH, L VH, R VH) from a given segmental level (C5, C6, C7, C8) in one model. COPE images from this stage were registered to PAM50 space using warp parameters generated during preprocessing (see main text).

Spatial extent of resting-state networks at group level was assessed using *randomise* (Winkler et al., 2014) with threshold-free cluster enhancement (5000 permutations, *p* < 0.003 (*p* = 0.05, Bonferroni corrected for 16 individual seed regions)).

#### 1.2 Results

Resting-state networks observed on MRI session 2 were highly similar to those observed on MRI session 1 (see Supplementary Figure 2 for overlap between clusters). Networks were largely confined to each segmental level, with sparser between-segment connections (Supplementary Figure 1 and 3). Similar to the results of session 1 data analysis, we observed a dorsal bias in functional connectivity of dorsal seeds and a ventral bias in functional connectivity of ventral seeds.

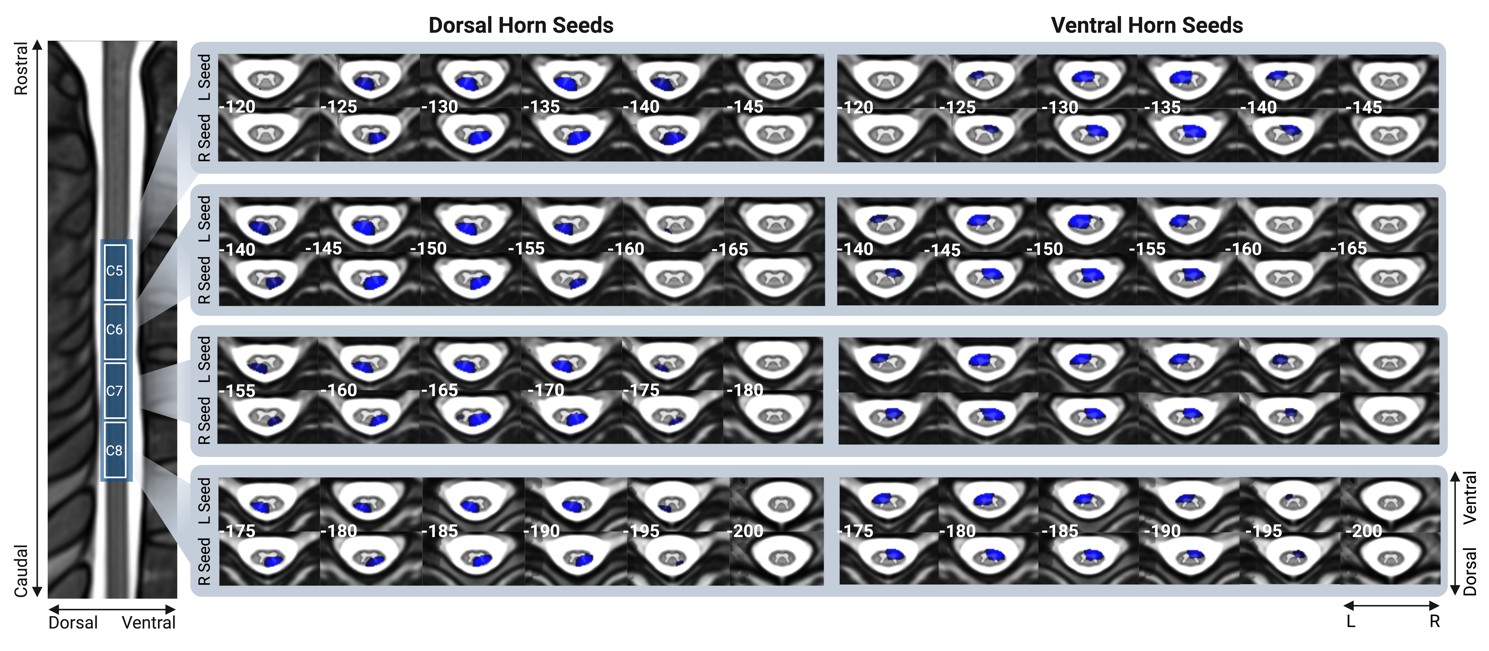

Supplementary Figure 1. Resting-state networks obtained from voxelwise connectivity analysis for each of the four quadrants (ventral/dorsal and left/right) of segmental levels C5-C8. Axial slices are marked with the *z* MNI coordinate. Each resting-state map was thresholded at *p* < 0.003 (*p* = 0.05, Bonferroni corrected for 16 individual seed regions).

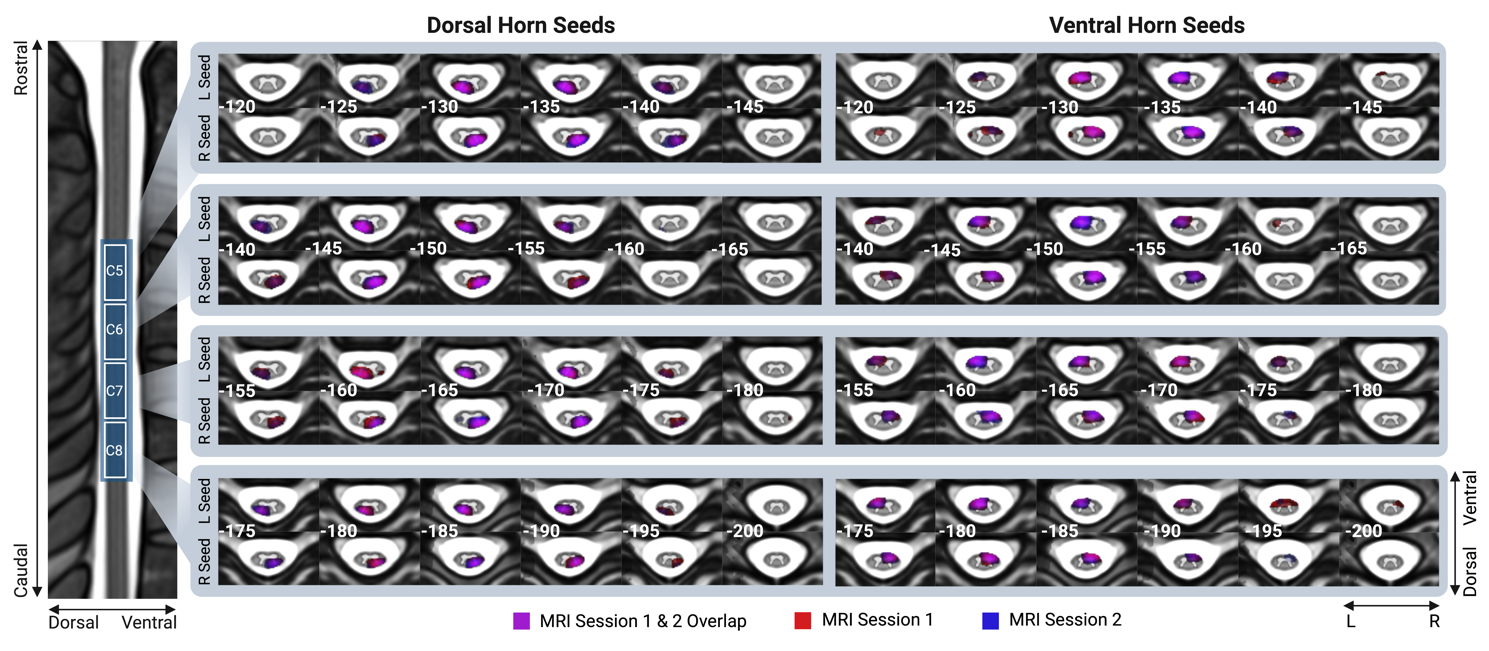

Supplementary Figure 2. Resting-state networks obtained from voxelwise connectivity analysis for each of the four quadrants (ventral/dorsal and left/right) of segmental levels C5-C8. Networks observed on MRI session 1 (red) and 2 (blue), as well as their overlap (purple) are displayed. Axial slices are marked with the *z* MNI coordinate. Each resting-state map was thresholded at *p* < 0.003 (*p* = 0.05, Bonferroni corrected for 16 individual seed regions).

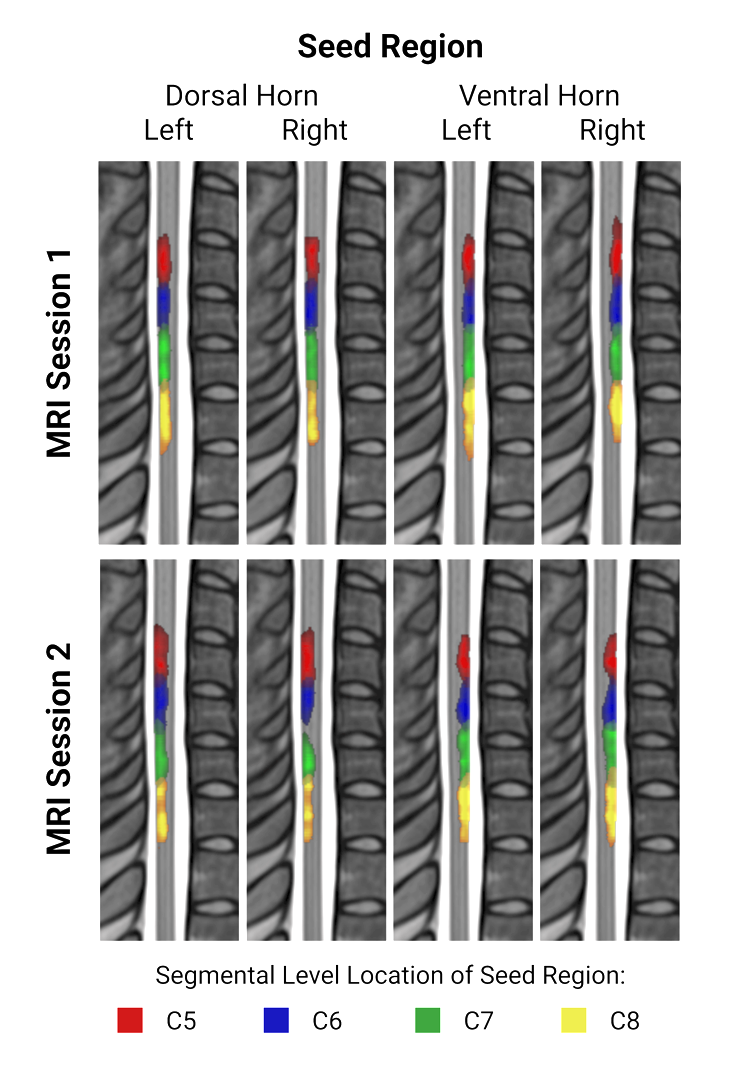

Supplementary Figure 3. A sagittal view (x = 0) of resting-state networks obtained from voxelwise connectivity analysis displaying sessions 1 (top row) and 2 (bottom row). Colours represent the location of seed regions used to estimate resting state networks (red = C5, blue = C6, green = C7, yellow = C8). Each resting-state map was thresholded at *p* < 0.003 (*p* = 0.05, Bonferroni corrected for 16 individual seed regions).

### 2. Seed-to-seed connectivity analysis for data acquired during MRI session 2

#### 1.3 Methods

Seed-to-seed connectivity analysis of data acquired during MRI session 2 was performed identically to that of MRI session 1 (described in the main text). Briefly, Pearson correlations were computed between each pair of seed regions at subject-level. The resultant correlation coefficients were *Z*-transformed. Statistical significance at group-level was assessed using a one-sample *t*-test. A positive false discovery rate (FDR) was used to account for multiple comparisons (thresholded at *p* < 0.05).

#### 1.4 Results

A correlation matrix depicting cervical spinal cord connections is shown in Supplementary Figure 3. Overall, the pattern of results was similar to that observed on session 1. On average, within segment, the strongest statistically significant positive correlations were observed within hemicord (i.e. ipsilateral DH-VH), followed by VH-VH and DH-DH connections, and DH-VH connections between hemicords (i.e. left DH – right VH, right DH – left VH). Weaker but statistically significant positive correlations were also observed between neighbouring segments, including DH-DH, VH-VH, as well as within and between hemicords. Finally, negative correlations were observed between the right VH of segment C8 and both left and right DH of segments C5 and C6.

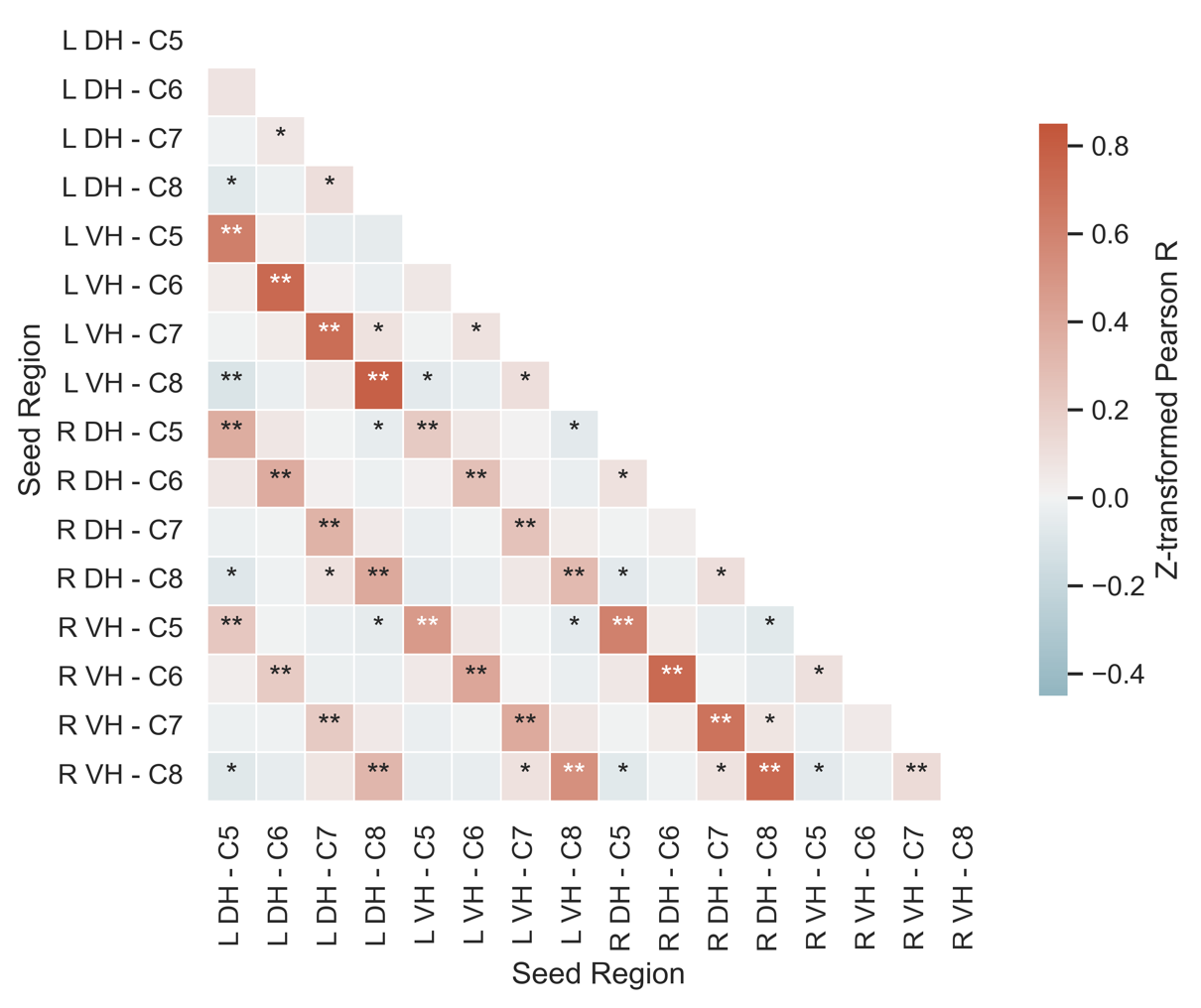

Supplementary Figure 4 Seed-to-seed correlation matrix displaying z-transformed Person R.

DH = Dorsal Horn, L = Left, VH = Ventral Horn, R = Right.

*p < 0.05, **p < 0.001
